# Iron availability regulates PIN-mediated auxin transport and distribution to modulate root gravitropic growth in *Arabidopsis*

**DOI:** 10.64898/2026.05.20.726447

**Authors:** Yao Fang, Mengjuan Kong, Yakun Peng, Shutang Tan

## Abstract

Iron (Fe) is an essential micronutrient for plant growth, and the hormone auxin is a key regulator of developmental processes, including root gravitropism. Here, we investigated the molecular mechanisms underlying the crosstalk between iron nutrition and auxin-mediated root growth in *Arabidopsis thaliana*. Phenotypic analysis revealed that iron deficiency strongly shaped root system architecture and root gravitropism, and these phenotypes were exacerbated in the iron uptake mutant *irt1-1*. Genetic analysis revealed that iron deficiency did not aggravate the gravitropic defect of the *pin2* mutant, *eir1-4*, suggesting that iron availability modulates root gravitropism through a *PIN2*-dependent pathway. Further transcriptomic analysis confirmed that iron deficiency significantly altered the expression of numerous genes related to the auxin pathway, providing molecular evidence for the observed physiological connection. Collectively, this study revealed that iron availability regulates root gravitropic growth by modulating PIN-mediated auxin transport and distribution, providing insights into how plants integrate nutritional cues with developmental programs.

**Graphical abstract**

**A brief description:** Iron modulates auxin transport and root tip distribution by regulating PIN2 protein, thereby mediating root gravitropism in *Arabidopsis*.

**Public summary:**

1. Iron nutrition specifically regulates root gravitropism and architecture in *Arabidopsis*.
2. Iron deficiency disrupts local auxin homeostasis in root tips and impairs asymmetric distribution in response to gravity.
3. Iron deficiency stress significantly reduces the abundance of PIN2 protein in root tip cells and disrupts its polar localization pattern on the plasma membrane, thereby precisely modulating polar auxin transport by interfering with the vesicle trafficking and recycling efficiency of PIN2.
4. RNA-seq results showed that iron deficiency induced differential expression of multiple auxin-related genes, indicating that iron nutrition affects root development through the auxin pathway.

## 1. Introduction

Iron (Fe) is an essential micronutrient for virtually all living organisms and plays a crucial role in various life activities (Kobayashi and Nishizawa, 2012). In plants, iron is indispensable for photosynthesis, cellular respiration and chlorophyll biosynthesis, making its homeostasis crucial for growth and development (Henriques et al., 2007; Hindt and Guerinot, 2012; Carvalhais et al., 2013; Romera et al., 2023). Although iron is abundant in the Earth’s crust, its bioavailability is often severely limited. In soils, iron predominantly exists as insoluble ferric (Fe^3+^) hydroxides, making it largely unavailable for plant uptake. Therefore, plants have evolved complex and tightly regulated strategies to acquire iron from the soil. In the model plant *Arabidopsis thaliana*, the primary strategy to acquire iron involves three key steps: acidification of the rhizosphere through proton release, reduction of Fe^3+^ into the soluble form (Fe^2+^) and subsequent uptake of Fe^2+^ into root cells (Kobayashi and Nishizawa, 2012). The final and critical uptake step is mediated by the plasma membrane transporter IRON-REGULATED TRANSPORTER 1 (IRT1) (Connolly et al., 2002; Henriques et al., 2007; Vert and Chory, 2009). The expression of *IRT1* is tightly controlled at both the transcriptional and posttranslational levels. IRT1 localizes at the polar plasma membrane (at outer lateral side of the epidermal cells) that requires SORTING NEXIN1 (SNX1) for correct trafficking (Barberon et al., 2014; Ivanov et al., 2014). For example, the abundance of *IRT1* is regulated by monoubiquitin-dependent endocytosis, which involves specific lysine residues and RING E3 ubiquitin ligases such as IRT1 DEGRADATION FACTOR1 (IDF1) (Barberon et al., 2011). Recent work supports the role of IRT1 as a transceptor (transporter and receptor), responsible for perceiving the intracellular Fe^2+^ levels, and regulating its own protein stability (Connolly et al., 2002; Dubeaux et al., 2018). The *irt1-1* mutant showed auxin defects, which was reported to be possibly related to the auxin transporter AUXIN1 (AUX1) (Giehl and Lima, 2012).

The plant hormone auxin acts as a central regulator of plant growth and development, controlling processes such as tropisms, organ patterning, and root system architecture (RSA) (Shen et al., 2015; Ogura et al., 2019; Tan et al., 2021; Roychoudhry et al., 2023; Luschnig and Friml, 2024). Auxin signalling regulates plant growth and development mainly through the transcriptional pathway mediated by the nuclear TRANSPORT INHIBITOR RESPONSE1 (TIR1)/AUXIN SIGNALLING F-BOX (AFB)-AUXIN/INDOLE-3-ACETIC ACID (Aux/IAA) coreceptor complex. This process releases the transcription factors Auxin Response Factors (ARFs), which controls the expression of downstream genes (Gray et al., 2001; Qi et al., 2022; Carrillo-carrasco et al., 2023; Kuhn et al., 2024; Chen et al., 2025). Recent research suggests this canonical pathway couples transcriptional regulation with a parallel cAMP production, which serves as a second messenger also required for downstream auxin responses (Qi et al., 2022; Chen et al., 2025). Additionally, the AUXIN BINDING PROTEIN1 (ABP1)/ ABP1 LIKEs (ABLs)-TRANSMEMBRANE KINASEs (TMKs) pathway perceives extracellular auxin and regulates many non-transcriptional cellular events (Li et al., 2021; Lin et al., 2021; Friml et al., 2022; Yu et al., 2023; Rodriguez et al., 2025; Huang et al., 2026). Auxin gradient establishment, which is crucial for its function, is dynamically mediated by polar auxin transport (PAT) through influx and efflux carriers, AUXIN1/LIKE AUXIN1s (AUX1/LAX) (Peret et al., 2012; Zhao et al., 2023; Yang et al., 2025) and the PIN-FORMED (PIN) family proteins (Petrášek et al., 2006; Tan et al., 2021; Yang et al., 2022; Pan et al., 2025; Zhang and Tan, 2026). The subcellular localization and abundance of these PIN proteins are highly regulated, thereby determining the direction of auxin flow (Wisniewska et al., 2006; Glanc et al., 2021; Tan et al., 2021; Konstantinova et al., 2022; Peng et al., 2024; Zhang and Tan, 2026).

Increasing evidence reveals the connection between iron availability and the auxin pathway in plants. For example, iron deficiency triggers auxin-mediated physiological responses and leads to shoot growth defects and photosynthesis depression in rice (Liu et al., 2015; Shen et al., 2015). In addition, exogenous IAA treatment increases Fe(III)-chelate reductase (FCR) activity in cucumber (Bacaicoa et al., 2011; Liu et al., 2015; Peng et al., 2024). These lines of evidence indicate that there is crosstalk between auxin signalling and iron deficiency signalling (Liu et al., 2015). Endogenous nitric oxide (NO) is a signalling molecule that regulates auxin transport proteins such as PIN2, and its production is related to both iron nutrition and the auxin response, thereby providing a mechanistic link (Romera et al., 2023). Notably, our recent research revealed that iron deficiency in *Arabidopsis* induces root wavy growth, a phenotype that may be related to the auxin response (Peng et al., 2024). However, the molecular mechanism underlying iron availability regulating root gravitropic growth is unclear.

In this study, we revealed that iron deficiency modulated auxin transport components, particularly PIN2 proteins, thereby interfering with the establishment of auxin gradients and ultimately disrupting root gravitropic responses. Understanding the interactions between metals and hormones is crucial for elucidating how plants adapt to fluctuating environments.

## 2 Materials and methods

### 2.1 Plant Materials and Growth Conditions

The *Arabidopsis thaliana* L. mutants and transgenic lines used in this study were generated in the Columbia-0 (Col-0) genetic background. The marker lines *pPIN1::PIN1-GFP* (Benková et al., 2003), *pPIN2::PIN2-Venus* (Leitner et al., 2012), *pPIN3::PIN3-GFP* (Zadnikova et al., 2010), *pPIN7::PIN7-GFP* (Zadnikova et al., 2010), *DR5rev::GFP* (Friml et al., 2003), the iron transporter mutant *irt1-1* (Connolly et al., 2002), the auxin-related mutants *arf7 arf19* (Okushima et al., 2007), *tir1-1 afb2-3* (Prigge et al., 2020), *pin3 pin4 pin7* (Zadnikova et al., 2010), *wei8-1* (Stepanova et al., 2008), and *eir1-4* (Abas et al., 2006) have been described previously. These reporter lines were introduced into the *irt1-1* background by crossing to generate *irt1-1 pPIN1::PIN1-GFP, irt1-1 pPIN2::PIN2-Venus, irt1-1 pPIN3::PIN3-GFP, irt1-1 pPIN7::PIN7-GFP*, and *irt1-1 DR5rev::GFP*. Table S1 lists all the plant lines analysed in this study, including mutants, reporter lines, and hybrid progeny.

For seedling phenotype analysis, seeds were surface sterilized with 75% ethanol and sown on 0.5× MS media (MES-buffered, pH 5.9; Kornet and Scheres, 2009) supplemented with 1% (w/v) sucrose and 0.8% (w/v) plant agar. The MS medium was subjected to different treatments, as detailed in Table S2. After stratification at 4°C for 2 days, the seeds were transferred to a 21°C plant growth chamber and grown vertically under long-day conditions (16 hours light/8 hours dark).

### 2.2 Root phenotype analysis

This study employed Col-0, the iron transport mutant *irt1-1*, and several auxin signalling-deficient mutants as materials. These samples were individually inoculated onto normal or specific element-deficient (Fe, Mn, Zn, Co) 1/2 MS solid media. Following two-day stratification at 4°C, all the materials were transferred to a greenhouse for vertical culture. Primary root length and lateral root number were assessed on days 7 and 11, respectively, to analyse the effects of nutrient deficiency (particularly iron deficiency) on root architecture and associated signalling pathways.

### 2.3 Root growth

The sterilized seeds were evenly sown in spots or scattered on 1/2 MS sucrose medium plates. The plates were sealed with parafilm and then placed in a refrigerator at 4°C for 2 days of stratification. The trays of cold-treated seeds were transferred to the greenhouse, where they were maintained at a constant temperature of 22°C under a photoperiod of 16 hours of light followed by 8 hours of darkness until seed germination and subsequent growth occurred.

### 2.4 Root gravitropic response

Four-day-old seedlings were subjected to a gravitropic experiment. The Petri dishes were rotated horizontally by 90° to apply gravitational stimulation. Representative images were captured at 0, 2, 4, 6, 8, 10, 12, and 24 hours post-treatment to document the progression of root bending. The root bending angles were quantitatively analysed via ImageJ software, with measurements based on the angle between the root tip direction and the gravitational vector direction. To systematically examine the changes in auxin distribution in root tips under gravistimulation, we employed the auxin-responsive reporter *DR5rev::GFP*. Four-day-old vertically grown wild-type Col-0 and *irt1-1* mutant seedlings were subjected to a 90° rotation, and after 4 hours, the spatial distribution of GFP signals was visualized using Confocal Laser Scanning Microscopy (CLSM).

### 2.5 Confocal laser scanning microscopy (CLSM) imaging

Confocal laser scanning microscopy (CLSM) imaging was performed via a Zeiss LSM 980 system equipped with a GaAsP detector (Zeiss, Germany). Images were acquired at 8-bit depth with 2× line averaging. The following imaging parameters were used for different fluorescent markers: for GFP-tagged proteins, 488 nm excitation and 495–545 nm emission were used; for Venus-tagged proteins, 514 nm excitation and 524–580 nm emission were applied. All subsequent image processing and analysis were conducted via Fiji software.

### 2.6 Pharmacological treatment with BFA and imaging

Wild-type Col-0 seedlings and *irt1-1* seedlings at the 4-day-old stage were treated with 50 μM BFA or MS medium (control) for one hour, followed by observation of *PIN2::PIN2-Venus* subcellular localization via confocal laser scanning microscopy.

### 2.7 Image Analysis and Morphological Analysis

To analyse the root morphology of the seedlings systematically, we first captured overall phenotypic images of the roots via a Sony A6000 camera equipped with a macrolens. The images were then quantitatively processed with ImageJ software, including measurements of primary root length, calculation of root tip curvature angles, and manual counting of lateral root numbers. The morphological characteristics of the root hairs were observed and imaged at high resolution via a stereomicroscope.

For fluorescence signal detection, fluorescence images of the root samples were acquired via a Zeiss LSM 980 laser scanning confocal microscope. All images were collected under consistent optical parameters. Subsequent quantitative analysis of fluorescence intensity in regions of interest, including fluorescence value statistics and distribution comparisons, was performed via ImageJ software to objectively evaluate differences in gene expression or signalling molecule localization.

### 2.8 RNA-seq analysis

Wild-type Arabidopsis thaliana Col-0 and the iron transport mutant *irt1-1* were cultured in iron-sufficient and iron-deficient media for 14 days. Plant samples of 100 mg were collected for transcriptome sequencing analysis. Library construction and sequencing were performed by BGI (Beijing Genomics Institute) using their proprietary BGISEQ high-throughput sequencing platform. The sequencing reads from each sample were aligned to the Arabidopsis thaliana reference genome (TAIR10 version), and data analysis was conducted using BGI’s Dr. Tom platform. Differentially expressed genes (DEGs) between iron-sufficient and iron-deficient conditions were screened using the thresholds of |log (fold change)| ≥ 1 and Q ≤ 0.05 (or P ≤ 0.05). For the identified DEGs, Gene Ontology (GO) biological process (GO_BP) enrichment analysis was performed. P-values were calculated using the phyper function in R language, and false discovery rate (FDR) correction was applied to obtain Q-values, with Q ≤0.05 as the threshold for significant enrichment. Gene expression heatmaps were generated using the OECloud online platform (https://cloud.oebiotech.com).

### 2.9 RT qPCR analysis

Reverse transcription quantitative PCR (RT qPCR) was used to detect the transcript levels of *PIN1, PIN2, PIN3, PIN4, PIN5, PIN6, PIN7*, and *PIN8* in Col-0 plants and *irt1-1* mutants grown on normal iron- and iron-deficient MS media. Additionally, the expression levels of *CEP5, JAZ10, ANT, MYB896, SAUR77, MYB31, GRDP2 and PIP2* in wild-type (Col-0) plants under normal iron- and iron-deficient conditions were analysed. *ACTIN7* (*AT5G09810*) was used as an internal reference gene for normalization.

Total RNA was extracted from plant tissues via the Trelief™ RNAprep Pure Tissue & Cell Kit (Tsingke, Cat# TSP 413) and treated with DNase to remove genomic DNA contamination. Reverse transcription was performed with 1 μg of RNA. The resulting cDNA was then amplified via SYBR Green qPCR Master Mix (Tolobio, Cat# 22204) on a Bio-Rad CFX Connect Real-Time PCR System. The relative expression levels of target genes were calculated via the 2^–ΔΔCt method, normalized to *ACTIN7* expression, and standardized by setting the expression level of wild-type plants under a specific tissue or condition as “1”. The experimental results are presented as the means ± standard deviations (SDs) of three biological replicates.

### 2.10 Quantification and statistics

Most experiments were repeated at least three times independently, with similar results obtained. To measure primary root length, photographs were analysed via ImageJ v1.53t (https://imagej.nih.gov/ij/download.html) (Schneider et al., 2012). The fluorescence intensity of the CLSM images was quantified via Fiji (https://fiji.sc/) (Schindelin et al., 2012). Data visualization and statistical analysis were mostly performed with GraphPad Prism 9.

### 2.11 Accession numbers

The sequence data from this study can be found in the *Arabidopsis* Genome Initiative or enBank/EMBL databases. The accession numbers are as follows: *IRT1* (AT4G19690), *CEP5* (AT5G66815), *JAZ10* (AT5G13220), *ANT* (AT4G37750), *MYB96* (AT5G62470), *SAUR77* (AT1G17345), *MYB31* (AT1G74650), *GRDP2* (AT4G37900), *PIP2* (AT437290), *PIN1* (AT1G73590), *PIN2* (AT5G57090), *PIN3* (AT1G70940), and *PIN7* (AT1G23080).

## 3 Results and Discussion

### 3.1 Iron deficiency induced by *irt1* mutation regulates seedling growth and development in *Arabidopsis*

To investigate the physiological effects of iron deficiency, we obtained a T-DNA insertion mutant, *irt1-1* (Connolly et al., 2002; Dubeaux et al., 2018), and explored the phenotypes of Col-0 and *irt1-1* seedlings grown on MS media supplemented with or without iron. We introduced the Gravitropism Index (GI) to precisely quantify the root response to gravity, calculated as the ratio of the actual length of the primary root to the straight-line distance between its two ends, thereby reflecting the degree of root curvature. Phenotypic analysis revealed that 7-day-old seedlings presented a more pronounced root wavy growth phenotype and a lower gravitropic index (distance/primary root length) (Peng et al., 2024) in the absence of iron (Fig. 1a-d). The *irt1-1* mutant presented a shorter primary root length and lower gravitropic index when grown on nutrient-sufficient media, and the root length and GI decreased further when iron was deficient (Fig. 1a-e). Notably, under iron-deficient conditions, *irt1-1* exhibited more severe growth defects, specifically manifested as a reduction in lateral root number, root hair number, and shorter root hairs (Fig. 1e-h). To evaluate the effects of other metal ions transported by IRT1 on the gravitropic response of plant roots, we conducted phenotypic observations under conditions of zinc deficiency, manganese deficiency, and cobalt deficiency (Fig. S1-S3). The results revealed that *irt1-1* did not exhibit severe defects in root growth or gravitropism, indicating that IRT1 functions primarily as an iron ion transporter.

**Fig. 1.**
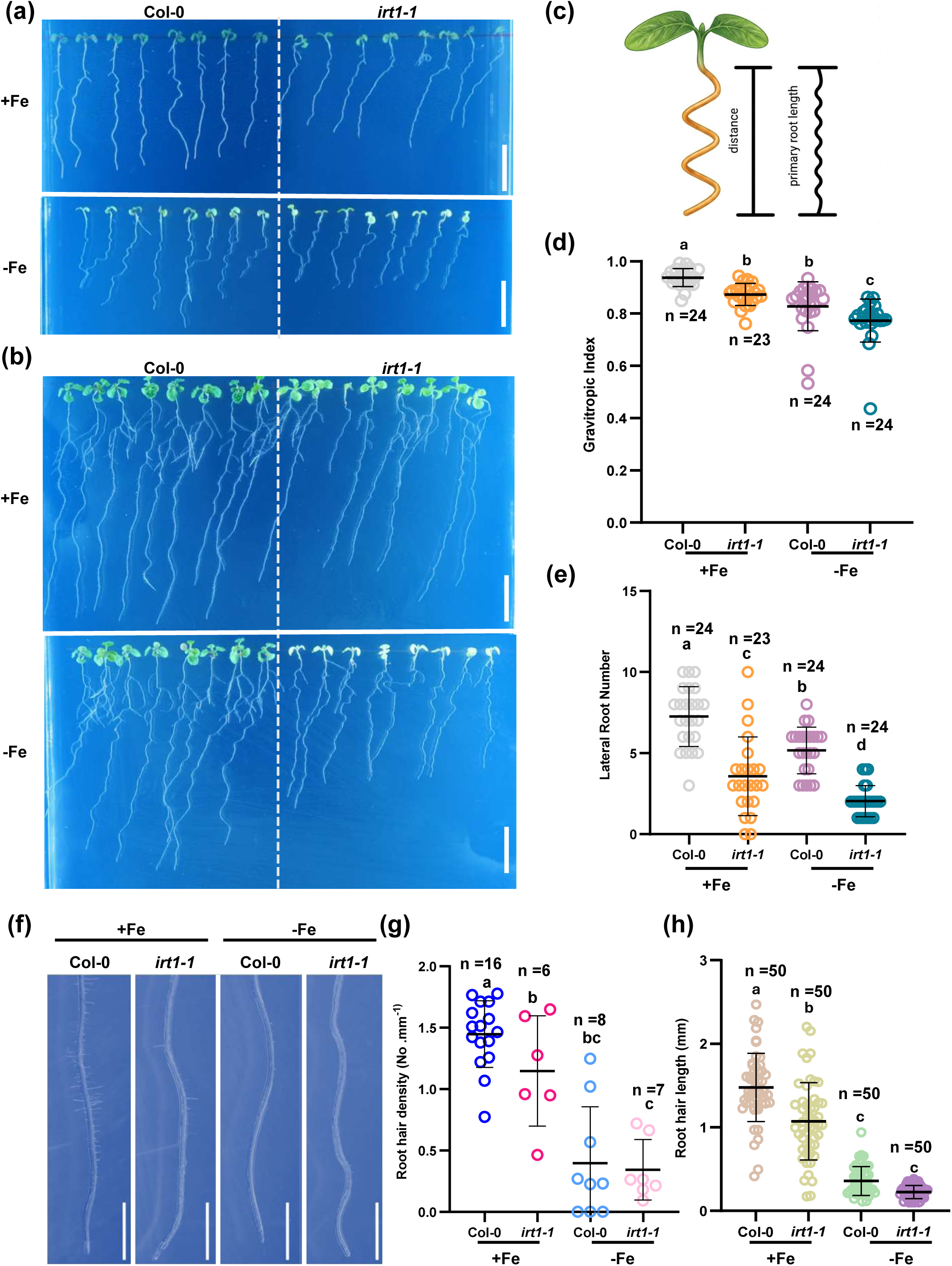
Iron specifically maintains the gravitropic growth of plants. (a, b) Phenotypic images of seven-day-old and eleven-day-old seedlings grown on the corresponding MS medium under normal iron and iron-deficient conditions. Scale bar, 1 cm. (c) Evaluation of the wavy root growth phenotype by measuring actual root length versus the straight-line distance between both ends. The root curvature is quantified by calculating the ratio of these values (i.e., the gravitropism index, GI). (d) *IRT1* deficiency resulted in a reduced GI, indicating a gravitropic defect. iron deficiency further decreased the GI, exacerbating the gravitropic defect. Measurements were taken using seven -day-old Col-0 and *irt1-1* seedlings under normal iron and iron-deficient conditions. The data are presented as the means ± SDs. Different letters denote significant differences; P < 0.05; one-way ANOVA with multiple comparisons. (e) Quantitative analysis of primary root length in *irt1-1* mutants compared with Col-0 under normal iron and iron-deficient conditions. *IRT1* deficiency resulted in shorter roots, with iron deficiency further exacerbating the defect. The data are presented as the means ± SDs. (f) *IRT1* mutation disrupts normal lateral root development. Lateral root number was measured in eleven -day-old Col-0 and *irt1-1* seedlings. The data are presented as the means ± SDs. Iron deficiency inhibited root hair development in Col-0, whereas *IRT1* deficiency results in nearly complete suppression of root hair growth. Root hair initiation was observed in seven-day-old seedlings under different iron conditions. The experiments were replicated eight times with consistent results. Scale bars, 2.5 mm. (g) Quantification of root hair density as shown. Data are presented as means ± SDs. Different letters indicate significant differences among groups by one-way ANOVA (P < 0.05). (h) The figure illustrates the root hair length of Col-0 and *irt1-1* under Fe-sufficient(+Fe) and Fe-deficient(-Fe) conditions. Each circle represents an individual measurement from independent biological replicates (n = 50). Data are presented as means ± SDs. Different letters indicate significant differences among groups by one-way ANOVA (P < 0.05).

To study the role of iron availability in the root gravitropic response in more detail, we inverted the vertically growing seedlings by 90° and measured the degree of root tip reorientation over time. This analysis revealed that *irt1-1* bent more slowly than did Col-0 with or without iron (Fig. 2a-c). These results suggest that iron deficiency caused by *IRT1* mutation may interfere with the auxin signalling pathway, leading to defects in root growth and the gravitropic response.

**Fig. 2.**
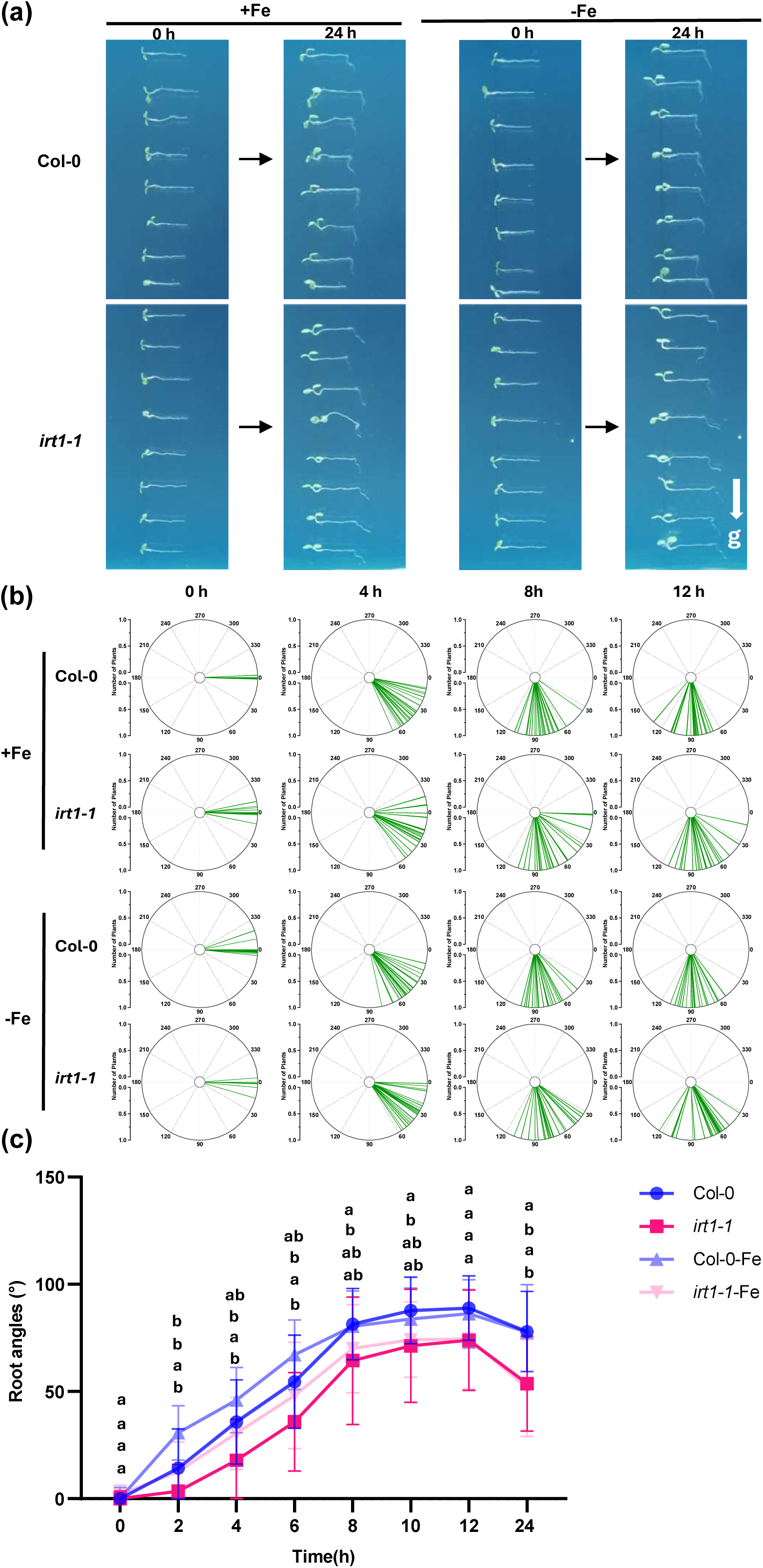
Iron deficiency delays the gravitropic response of Arabidopsis thaliana. (a) Representative images at 0 h and 24 h after the medium was rotated to 90°, followed by 4 days of growth on iron-sufficient (+Fe) MS media and iron-deficient (-Fe) MS media for wild-type (Col-0) seedlings and the *irt1-1* mutant. Scale bars, 1 cm. (b) The Root tip angles were measured at 0 h, 4 h, 8 h, and 12 h after gravitropic stimulation, four-day-old Col-0 and *irt1-1* seedlings were grown under iron-sufficient (+Fe) and iron-deficient (-Fe) conditions., presented as polar histograms. (c) Quantitative analysis of the root bending angles. At 2 h, 4 h, 6 h, 8 h, 10 h, 12 h, and 24 h, the root bending angles of the *irt1-1* mutant were significantly smaller than those of Col-0. The *irt1-1* mutant (*irt1-1*-Fe) exhibited impaired gravitropic responses, whereas Col-0 similarly displayed impaired gravitropic responses under iron deficiency. The data are presented as the means ± SDs. Different letters indicate significant differences among groups by one-way ANOVA (P < 0.05).

### 3.2 Iron homeostasis modulates auxin-mediated growth pathways

Auxin plays an essential role in regulating plant growth and gravitropism (Bennett et al., 1996; Tan et al., 2020; Tan et al., 2021; Xia et al., 2023; Wang et al., 2024). The auxin efflux carrier PIN family is essential for controlling auxin polar transport to maintain plant growth and development (Tan et al., 2021; Yang et al., 2022; Luschnig and Friml, 2024). To explore the role of auxin in iron-modulated root growth, we treated Col-0 and *eir1-4* (also named *pin2-T*) plants under normal iron- and iron-deficient conditions. The root phenotype indicated that iron deficiency did not aggravate the gravitropic defect of *eir1-4* (Fig. 3a-d and Fig. S4a-c), but it still reduced the number of lateral roots and root hairs, as well as the length of root hairs, in *eir1-4*. (Fig. 3e-h). However, further phenotype analysis of *pin347* revealed that *pin347* was still sensitive to iron deficiency in terms of root growth, development and the gravitropic response (Fig. S5 and 6). Given the essential roles of PIN2 in regulating auxin polar transport and root gravitropism, we speculate that the auxin pathway might be disrupted when iron availability is inhibited.

**Fig. 3.**
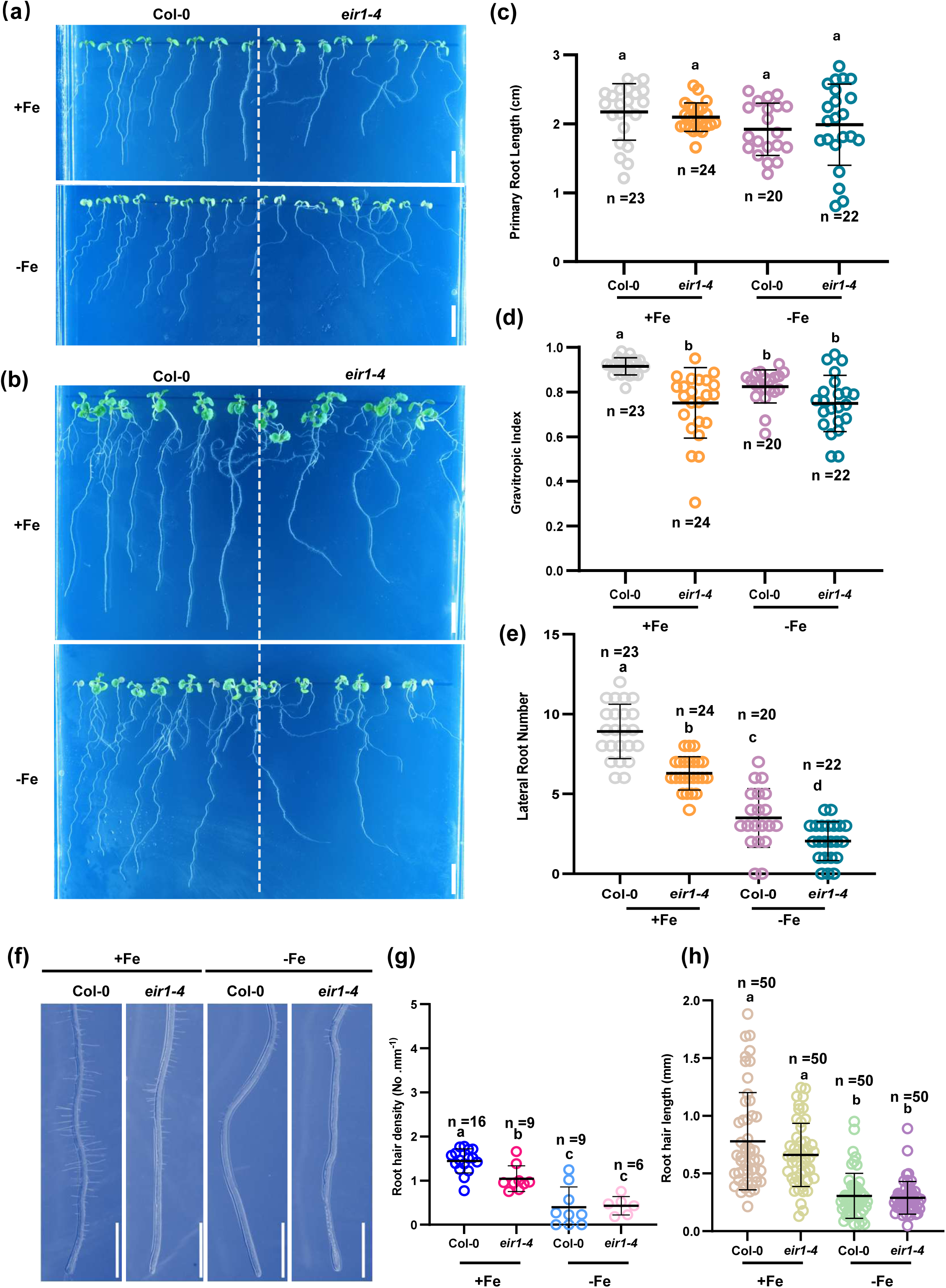
The gravitropism index of *eir1-4* showed no significant change under iron-deficient condition. **(a, b)** Representative images of seven-day-old and eleven-day-old Col-0 and *eir1-4* seedlings grown on corresponding MS media under normal iron- and iron-deficient conditions. Scale bars, 1 cm. (c) Quantitative analysis of primary root length in *eir1-4* mutants compared with Col-0 under normal iron and iron-deficient conditions. *PIN2* deficiency did not affect primary root length. The data are presented as the means ± SDs. (d) The absence of *PIN2* reduced GI, resulting in a gravitropic defect, whereas iron deficiency did not affect its GI. Measurements were taken using seven-day-old seedlings under both normal iron and iron-deficient conditions. The data are presented as the means ± SDs. (e) *PIN2* mutation disrupts normal lateral root development. Lateral root numbers were measured in eleven-day-old Col-0 and *eir1-4* seedlings. The data are presented as the means ± SDs. (f) Iron deficiency inhibited root hair development in wild-type (Col-0) roots, whereas *PIN2* deficiency severely suppressed root hair growth. The observation of root hair initiation in seven-day-old Col-0 and *eir1-4* seedlings was performed in eight replicates, yielding consistent results. Scale bars, 2.5 mm. (g) Quantification of root hair density as shown. Data are presented as means ± SDs. Different letters indicate significant differences among groups by one-way ANOVA (P < 0.05). (h) The figure illustrates the root hair length of Col-0 and *eir1-4* under Fe-sufficient(+Fe) and Fe-deficient(-Fe) conditions. Each circle represents an individual measurement from independent biological replicates (n = 50). Data are presented as means ± SDs. Different letters indicate significant differences among groups by one-way ANOVA (P < 0.05).

### 3.3 Iron deficiency interferes with PIN-dependent auxin transport

To verify this hypothesis, we introduced the auxin-responsive reporter *DR5rev::GFP* (Friml et al., 2003) into *irt1-1* by crossing. Quantitative confocal laser scanning microscopy (CLSM) analysis revealed a decreased auxin signal intensity in *irt1-1* root tips, and the signal intensity further increased under iron deficiency conditions (Fig. 4a and b). Further analysis of the *DR5rev::GFP* reporter in response to root gravitropism upon 90° reorientation revealed that auxin redistribution was reduced in *irt1-1*, especially under iron deficiency conditions (Fig. 4c and d). In addition, we tested the sensitivity of TIR1/AFB-Aux/IAA-ARF-related mutants to iron deficiency. In terms of the effects on gravitropism, the GIs of *wei8-1* and *tir1-1 afb2-3* were reduced, similar to those of Col-0, when iron was deficient (Fig. S7a-d and Fig. S8a-d). Further physiological experiments revealed that iron deficiency did not lead to a further reduction in lateral root number or root hair formation, which is inherently less common in these mutants. (Fig. S7e-h and Fig. S8e-h). Notably, *arf7 arf19* was insensitive to iron deficiency, as a lack of iron did not cause a reduction in the GI or changes in the number of root hairs or lateral roots (Fig. S9a-h). The results of the experiment involving a 90° rotation of the gravity direction also demonstrated that iron deficiency did not affect the rate of bending toward gravity in *arf7 arf19* (Fig. S10). These results indicate that the gravitropism defect caused by iron deficiency in plants is the result of disrupted auxin transport and distribution, as well as the inhibition of ARF-mediated auxin signalling.

**Fig. 4.**
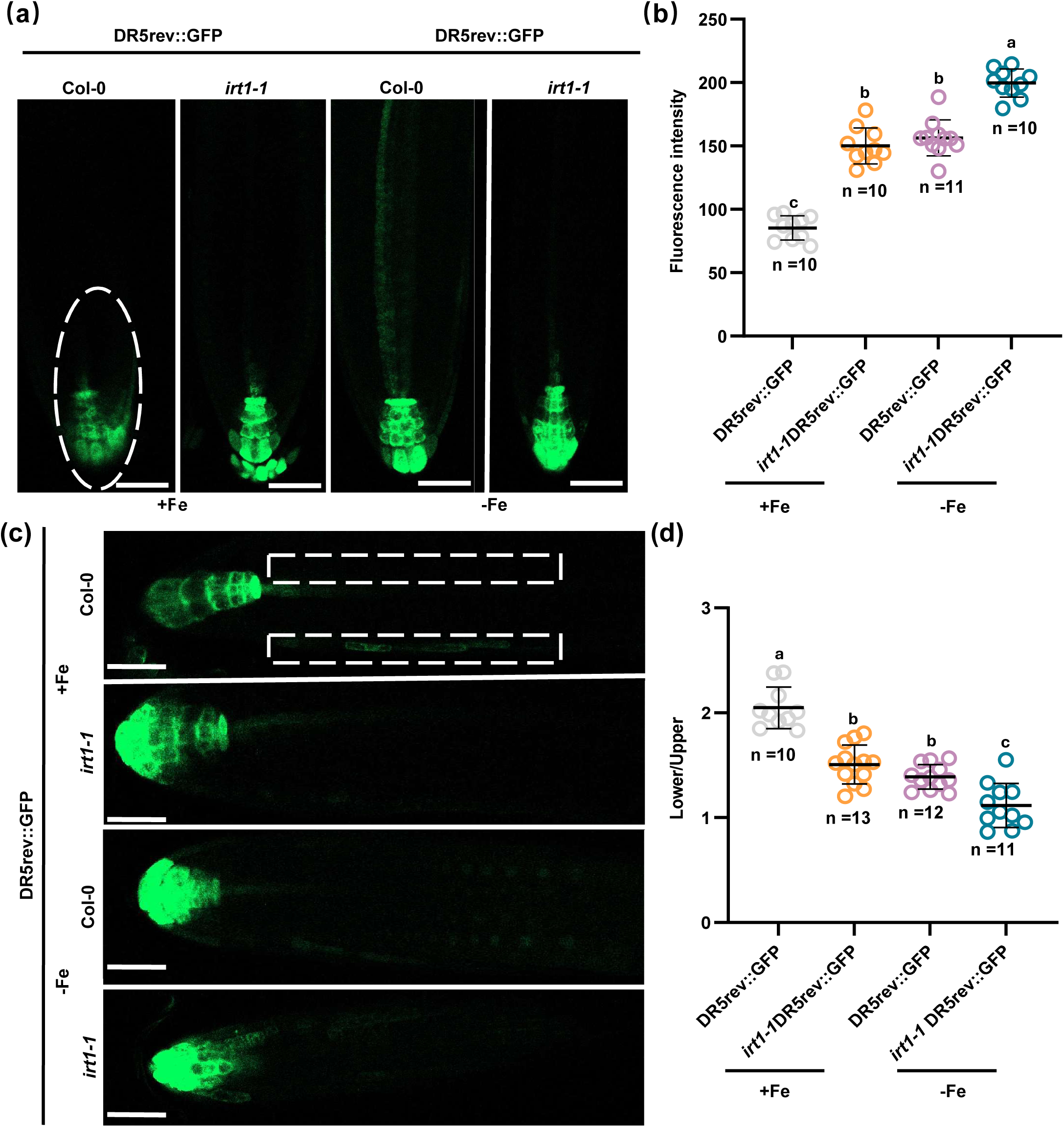
Iron deficiency affects the distribution and relocalization of auxin in root tips. (a) Images of *DR5rev::GFP* under different iron conditions in *Col-0* and *irt1-1*. The fluorescence intensity of *DR5 rev::GFP* was measured in five-day-old Col-0 seedlings and *irt1-1* seedlings. All samples were imaged from an identical root tip region. Laser wavelength, 488 nm, 5.00%; 0.0% offset; detector gain, 600 V. Scale bar, 20 μm. (b) Both iron deficiency and *IRT1* deficiency increased *DR5rev::GFP* fluorescence. The fluorescence intensity was measured in regions of equal size to characterise the auxin content via reporter expression. The data are presented as the means ± SDs; different letters denote significant differences; P < 0.05; one-way ANOVA for multiple comparisons. The experiments were independently replicated three times with similar results. (c) Representative images of *DR5rev::GFP* and *irt1-1*×*DR5rev::GFP* after 4 hours of gravity stimulation treatment under normal iron and iron-deficient conditions. Col-0 exhibited defects in gravity-stimulated *DR5rev::GFP* relocalisation under iron deficiency, whereas *irt1-1* plants also displayed defects in gravity-stimulated DR5rev::GFP relocalisation. Laser wavelength, 488 nm; 5.00%; 0.0% offset; detector gain, 600 V. Scale bar, 20 μm. (d) Compared with Col-0, *irt1-1* presented defects in gravity-stimulated *DR5rev::GFP* relocalisation, which was exacerbated by iron deficiency. The fluorescence at the base and apex was measured within fixed localised areas, and ratios were calculated. The data are presented as the means ± SDs; different letters denote significant differences; P < 0.05; one-way ANOVA for multiple comparisons; dots represent individual plants. The experiments were independently replicated three times with similar results.

To explore the role of PINs in this process, we introduced *PIN1::PIN1-GFP PIN2::PIN2-Venus, PIN3::PIN3-GFP* and *PIN7::PIN7-GFP* reporters into the *irt1-1* background by genetic crossing. CLSM revealed that the signal intensity of PIN2 was weakened in Col-0 plants under iron-deficient conditions and further diminished when PIN2 was introduced into the *irt1-1* background (Fig. 5a and b). Notably, analysis of PIN2 signal distribution in epidermal cells revealed that the polar localization of PIN2 on the plasma membrane was inhibited under iron-deficient conditions and that mutation of IRT1 exacerbated this defect (Fig. 5c and d). Brefeldin A (BFA) inhibits the process of PIN2 targeting from endosomes to the plasma membrane by disrupting protein cycling between the plasma membrane and the endomembrane system. There were fewer and smaller BFA bodies in *irt1-1* after BFA treatment, and this phenomenon was exacerbated under iron deficiency conditions, suggesting that the trafficking of PIN2 cargo was affected (Fig. 6a-d). Under iron-sufficient conditions, the *PIN1::PIN1-GFP* fluorescence signal in Col-0 was relatively weak, whereas iron deficiency significantly induced its enhancement. In contrast, the PIN1 signal in the *irt1-1* mutant was insensitive to fluctuations in iron levels. Under normal iron conditions, the PIN1 signal already exhibited an enhanced expression similar to that observed under iron deficiency, and no further upregulation was detected upon iron deficiency treatment, suggesting that loss of *IRT1* might cause the plant to be in a constitutive state of auxin transport disruption. The expression of *PIN7::PIN7-GFP* also showed a marked dependence on iron signals. Under iron deficiency, the *PIN7::PIN7-GFP* fluorescence intensity in Col-0 plants was significantly decreased. In *irt1-1* mutants, the *PIN7::PIN7-GFP* signal intensity under iron-sufficient conditions was already significantly lower than that in the wild-type control, and iron deficiency did not further alter the fluorescence. These cytological results indicated that iron deficiency affected the cellular abundance of the auxin polar transporters PIN1 and PIN7. In stark contrast to PIN1 and PIN7, the expression of *PIN3::PIN3-GFP* was not affected by iron signals. Under both iron-sufficient and iron-deficient conditions, there were no significant differences in the signal intensity or distribution pattern of *PIN3::PIN3-GFP* between Col-0 and *irt1-1* mutants. (Fig. S11). These results suggest that iron signals might regulate root development under iron-deficient conditions by selectively modulating the expression and localization of specific PIN proteins, thereby affecting auxin distribution.

**Fig. 5.**
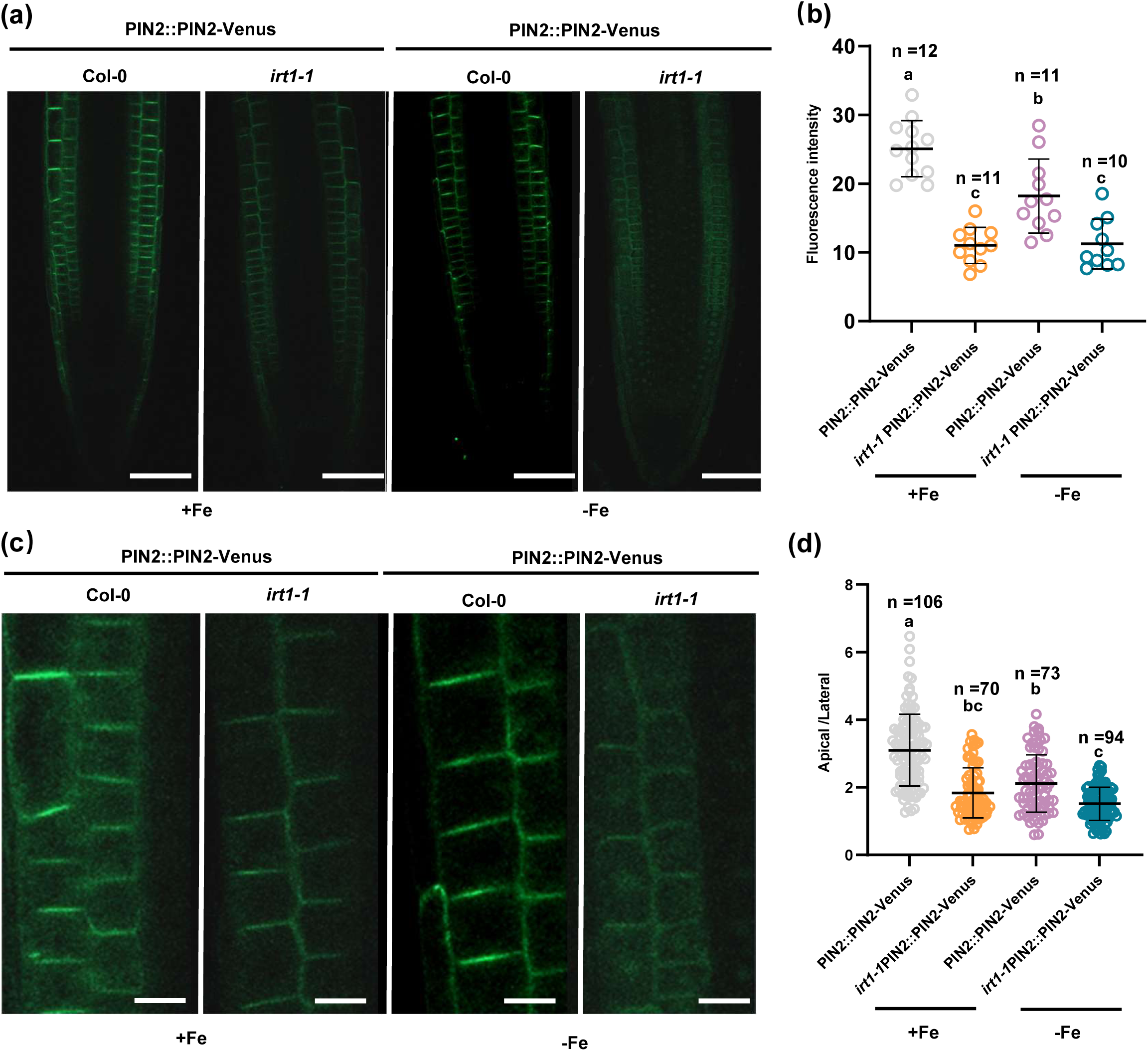
Iron deficiency interferes with the localization and polarity of PIN2 protein. (a) presentative images of *PIN2::PIN2-Venus* expression in the epidermal cells of *irt1-1* seedlings grown for 5 days on normal iron and iron-deficient MS media, with Col-0 as a control. Laser wavelength, 488 nm; 5.00%; 0.0% offset; detector gain, 650 V. Scale bar, 50 μm. (b) Compared with Col-0, *PIN2::PIN2-Venus* expression was lower in the epidermal cells of *irt1-1* seedlings, and iron deficiency exacerbated this defect. The fluorescence intensity was measured in regions of equal size to visualise reporter expression. The data are presented as the means ± SDs; different letters denote significant differences, P < 0.05; one-way ANOVA for multiple comparisons. The experiments were independently replicated three times with similar results. (c) Normal iron uptake (mediated by the IRT1 protein) was essential for maintaining the correct polarised localisation of the PIN2 protein on the cell membrane. Subcellular localisation of *PIN2::PIN2-Venus* in the epidermal cells of *irt1-1* seedlings under normal iron and iron-deficient conditions. Under normal iron conditions, the polarised localisation of *PIN2::PIN2-Venus* in *irt1-1* is partially disrupted, with reduced signalling at the plasma membrane. Under iron deficiency, the polarised localisation of *PIN2::PIN2-Venus in irt1-1* was completely lost. This protein forms distinct intracellular aggregates dispersed in the cytoplasm. In contrast, *PIN2::PIN2-Venus* in Col-0 clearly localised to the cell membrane (polarised localisation) regardless of iron availability. The fluorescence intensity at the same magnification was shown to indicate reporter gene expression. Scale bar, 5 μm. (d) Polarised localisation of PIN2 was indicated by the ratio of apical to lateral fluorescence signals. The data are presented as the means ± SDs; These data were derived from 101, 68, 64, and 88 cells from 10 seedlings. Different letters denote significant differences; P < 0.05; one-way ANOVA with multiple comparisons.

**Fig. 6.**
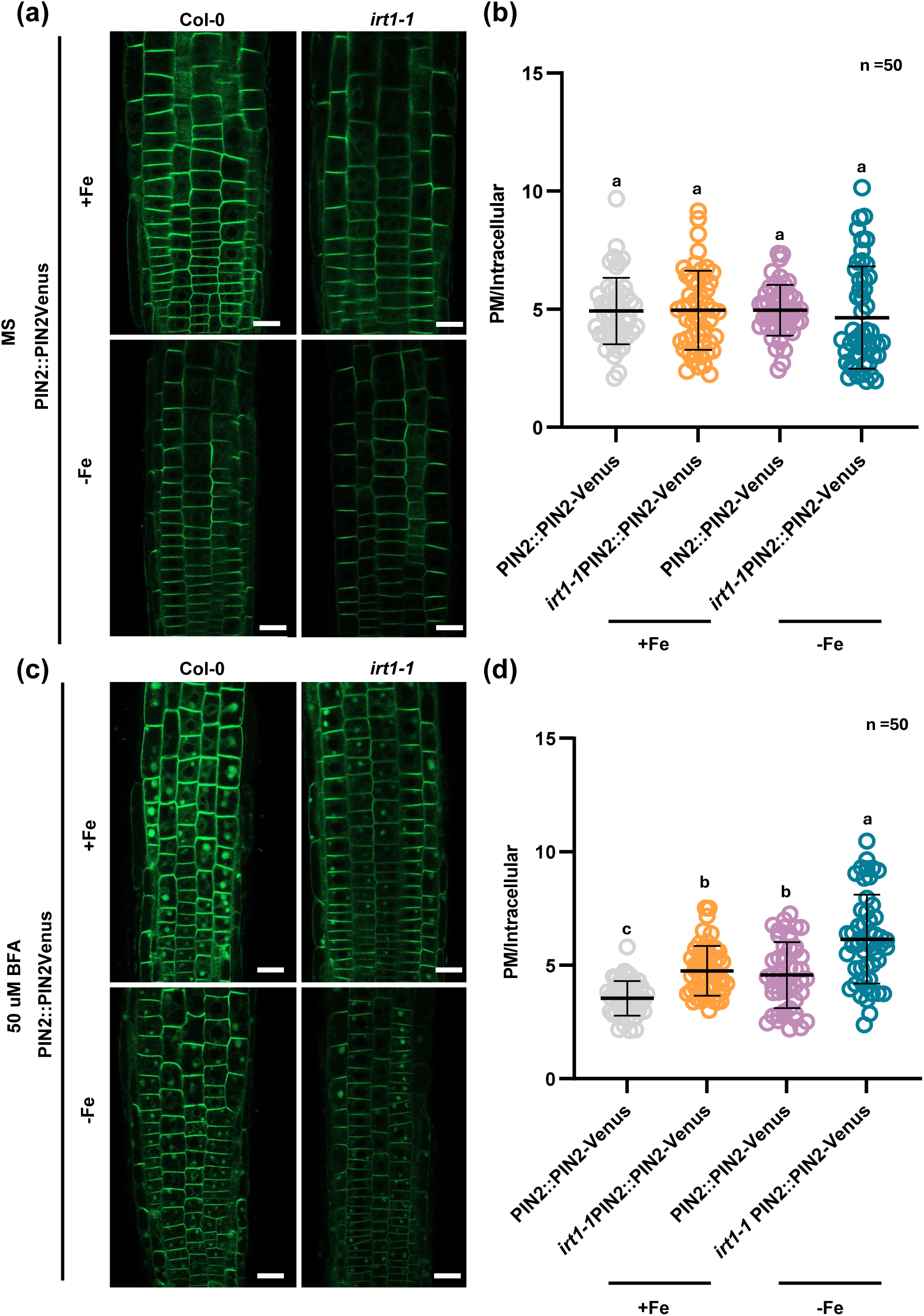
Iron deficiency impairs the recycling and vesicular trafficking of PIN2 protein. (a) Confocal images showing *PIN2::PIN2-Venus* localisation in four-day-old Col-0 and *irt1-1* seedlings after 1 hour of treatment with MS mixture. Laser wavelength: 488 nm, 5.00%; detector gain: 600 V. Scale bar, 20 μm. (b, d) Quantitative analysis of *PIN2::PIN2-Venus* signals on the plasma membrane of basal epidermal cells: the proportion of intracellular signals. Experimental data are presented as the mean ± SDs, with the sample size n indicated in the figure; different letters indicate significant differences (P<0.05), and post-hoc tests were performed following one-way analysis of variance (ANOVA). This experiment was independently replicated three times with consistent results. (c) Confocal images showing *PIN2::PIN2-Venus* localisation in four-day-old Col-0 and *irt1-1* seedlings after 1 hour of treatment with 100 μM BFA. Laser wavelength: 488 nm, 5.00%; detector gain: 600 V. Scale bar, 20 μm.

### 3.4 Iron deficiency remodels the auxin-related transcriptome in *Arabidopsis*

To further investigate the molecular mechanisms underlying the effects of iron deficiency, we performed RNA sequencing (RNA-seq) profiling experiments with Col-0 and *irt1-1* seedlings grown on MS media supplemented with or without iron. Through analysis of the results, we discovered a significant number of differentially expressed genes (DEGs) (Fig. S12). Further GO analysis showed that under iron deficiency, most of these differentially expressed genes (DEGs) were enriched in pathways related to defense responses and metabolic processes (Fig. 7a and Fig. S13). The heatmap of auxin-related DEGs indicated that iron deficiency also impacts the auxin signalling pathway (Fig. 7b). To determine the effects of iron deficiency on the auxin signalling pathway, we conducted RT qPCR analysis of these DEGs. The expression levels of *CEP5, JAZ10, ANT* and *MYB96* decreased under iron-deficient conditions, and the expression levels of *SAUR77, MYB31, GRDP2* and *PIP2* increased (Fig. 7c). JA regulates root auxin distribution by influencing auxin biosynthesis and transport. As inhibitors of jasmonic acid, *JAZ* proteins downregulate *JAZ10* under iron stress, disrupting the balance between jasmonic acid and auxin signalling (Kazan and Manners, 2009; Westfall et al., 2012). This leads to disordered auxin distribution, thereby inhibiting root geotropism. The expression of *CEP5* is negatively regulated by auxin. Iron deficiency affects auxin distribution or signalling, thereby downregulating *CEP5*. The *AINTEGUMENTA* (*ANT*) transcription factor is a key regulator of auxin synthesis that promotes auxin production by directly activating the *YUCCA4* gene (Vernoux et al., 2000). Under iron deficiency stress, *ANT* expression is suppressed, directly leading to reduced transcriptional levels of its downstream target gene *YUCCA4* and consequently causing a reduction in localized auxin synthesis. *MYB96* serves as a crucial inhibitor of callus formation. The *MYB94/96*-*LBD29* pathway, which acts as a novel regulatory mechanism, operates in parallel with the ARF7/19-LBD pathway (Okushima et al., 2007; Dai et al., 2020). Environmental stress (iron deficiency) reshapes the hormonal signalling network within plants by modulating the key transcription factor *MYB96*, thereby increasing the inhibition of callus formation to facilitate adaptation to stress. *GRDP2* affects auxin distribution by regulating the subcellular localization of PIN family proteins, which are key carriers for polar auxin transport. In *grdp2 Arabidopsis* mutants, PIN1, PIN2, and PIN7 are abnormally retained in the cytoplasm instead of the cell membrane, resulting in impaired auxin transport. Under iron deficiency stress, the expression of *GRDP2* is upregulated, which enhances the proper localization of PIN proteins and promotes auxin transport to adapt to stress. The upregulation of *SAUR77* under iron deficiency is closely associated with auxin. The SAUR gene family is composed of auxin responsive genes (Spartz et al., 2014; Du et al., 2020). Under iron deficiency, plants exhibit alterations in auxin synthesis, transport, and signal transduction, which in turn induce the upregulation of *SAUR77* gene expression. According to the literature, *PIP2* is an auxin-responsive gene. Iron deficiency leads to an increase in auxin, which in turn upregulates *PIP2* (Wang et al., 2021). *MYB* transcription factors are involved in a variety of secondary metabolic processes and stress responses. The upregulation of *MYB31* may affect lignin deposition or induce the production of other metabolites by regulating pathways such as phenylpropanoid metabolism. This, in turn, alters the characteristics of cell walls and indirectly influences auxin-mediated root system architecture (RSA). These results were consistent with the RNA-seq results, confirming that iron deficiency affects the auxin signalling pathway.

**Fig. 7.**
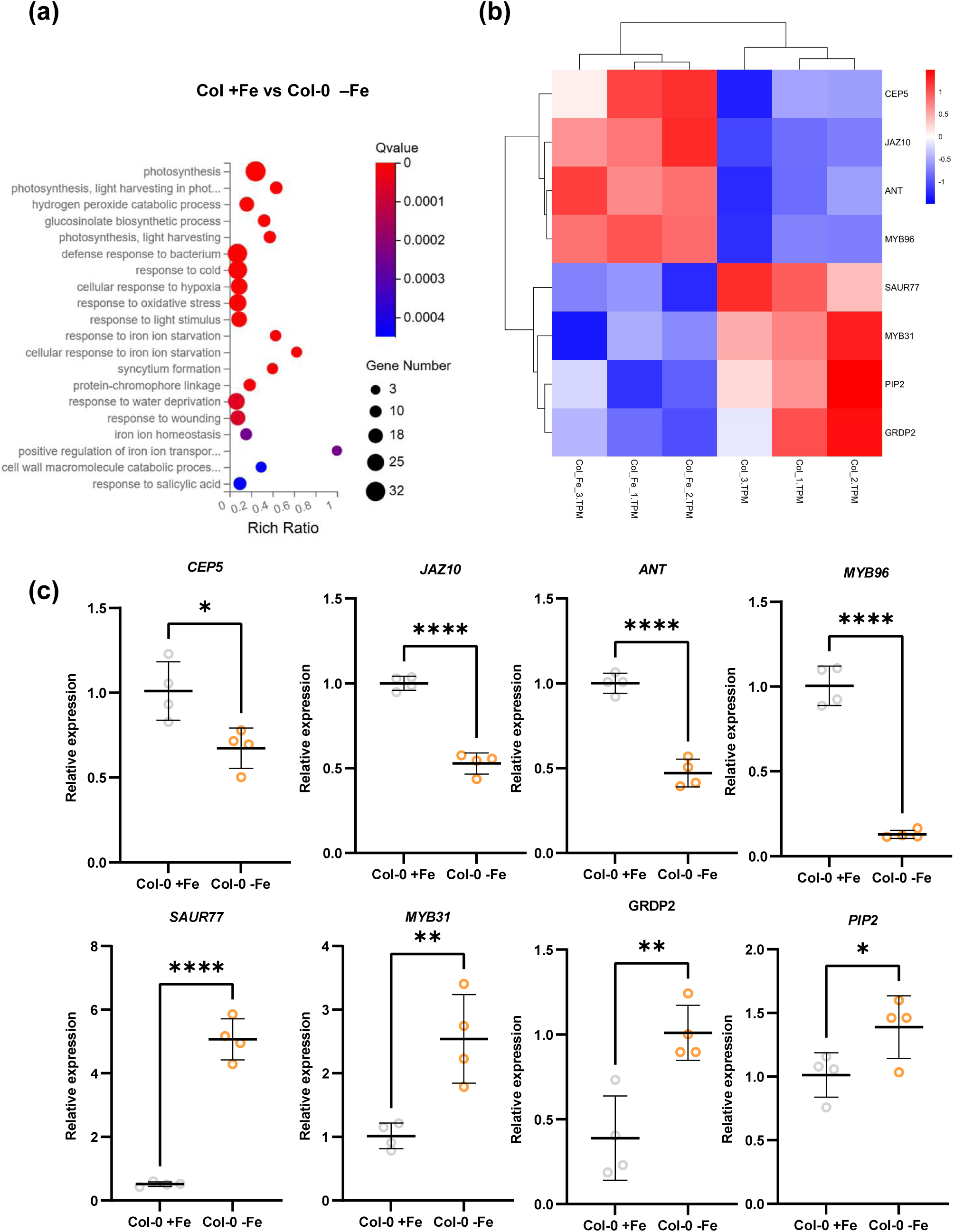
RNA-seq analysis indicating that iron signalling is associated with the auxin pathway. (a) Statistical results of the GO enrichment analysis of genes significantly differentially expressed in Col-0 under normal iron and iron-deficient conditions. (b) Heatmap displaying differentially expressed genes in Col-0 under normal iron and iron-deficient conditions. (c) Real-time quantitative PCR analysis of the relative expression levels of the auxin-related genes *CEP5, JAZ10, ANT, MYB896, SAUR77, MYB31, GRDP2*, and *PIP2*. The dots represent individual data points (Col-0+Fe: silver dots; Col-0+Fe: yellow dots). The solid lines denote the means ± SDs. *p<0.05, **p<0.01, ***p<0.001, and ****p<0.0001 (Welch’s t test).

In addition, we further analyzed the transcript levels of *PIN1* to *PIN8* under iron-deficient conditions (Fig. S14). The results showed that iron signals exhibited a selective and differential regulation among PIN family members. In the *irt1-1* mutant subjected to iron deficiency stress (*irt1-1*-Fe), the transcript levels of *PIN1, PIN2, PIN5*, and *PIN8* were significantly downregulated compared with those under normal iron conditions. Consistent with the aforementioned protein localization experiments, the transcriptional downregulation of *PIN1* was in agreement with its reduced protein abundance, indicating that iron signals tightly control *PIN1* expression at both the transcriptional and translational levels. For *PIN2*, although its transcript levels remained relatively stable in the Col-0, they were still transcriptionally repressed in *irt1-1* under severe iron deficiency. In stark contrast to the aforementioned genes, *PIN3, PIN4, PIN6*, and *PIN7* exhibited significant transcriptional upregulation under iron-deficient conditions in the *irt1-1* mutant. Notably, for *PIN3* and *PIN7*, their transcript abundances dramatically increased in the roots of *irt1-1* under iron deficiency. Collectively, these results indicated that iron signals do not simply inhibit auxin transport but rather act through fine-tuned transcriptional reprogramming: on one hand, iron signal downregulated PIN proteins (PIN1/PIN2) responsible for primary polar auxin transport to suppress primary root elongation, on the other hand, iron signal upregulated the transcription of PIN proteins (PIN3/PIN4/PIN7) involved in lateral/gravitropic responses, attempting to remodel root system architecture in response to the stress.

## 4 Discussion

Iron (Fe) is an essential micronutrient for plant growth and development (Kobayashi and Nishizawa, 2012). In this study, our results reveal that iron plays an essential role in regulating root gravitropic growth, largely replying on modulating the auxin distribution and the subsequent signalling pathway. Phenotypic analysis revealed that iron deficiency strongly shaped root system architecture and root gravitropism, and these phenotypes were exacerbated in the iron uptake mutant *irt1-1*. Genetic analysis revealed that iron deficiency did not aggravate the gravitropic defect of the *pin2* mutant, *eir1-4*, suggesting that iron availability modulates root gravitropism through a *PIN2*-dependent pathway. Notably, the decrease in iron levels impair the polar PM localization of PIN2 protein, suggesting a potential regulatory mechanism involving the endomembrane trafficking of PIN2 (Fig. 8), for which the underling mechanism awaits further investigation. We speculate that there might be an iron-binding or -responsive protein(s) involved in the proposed pathway.

**Fig. 8.**
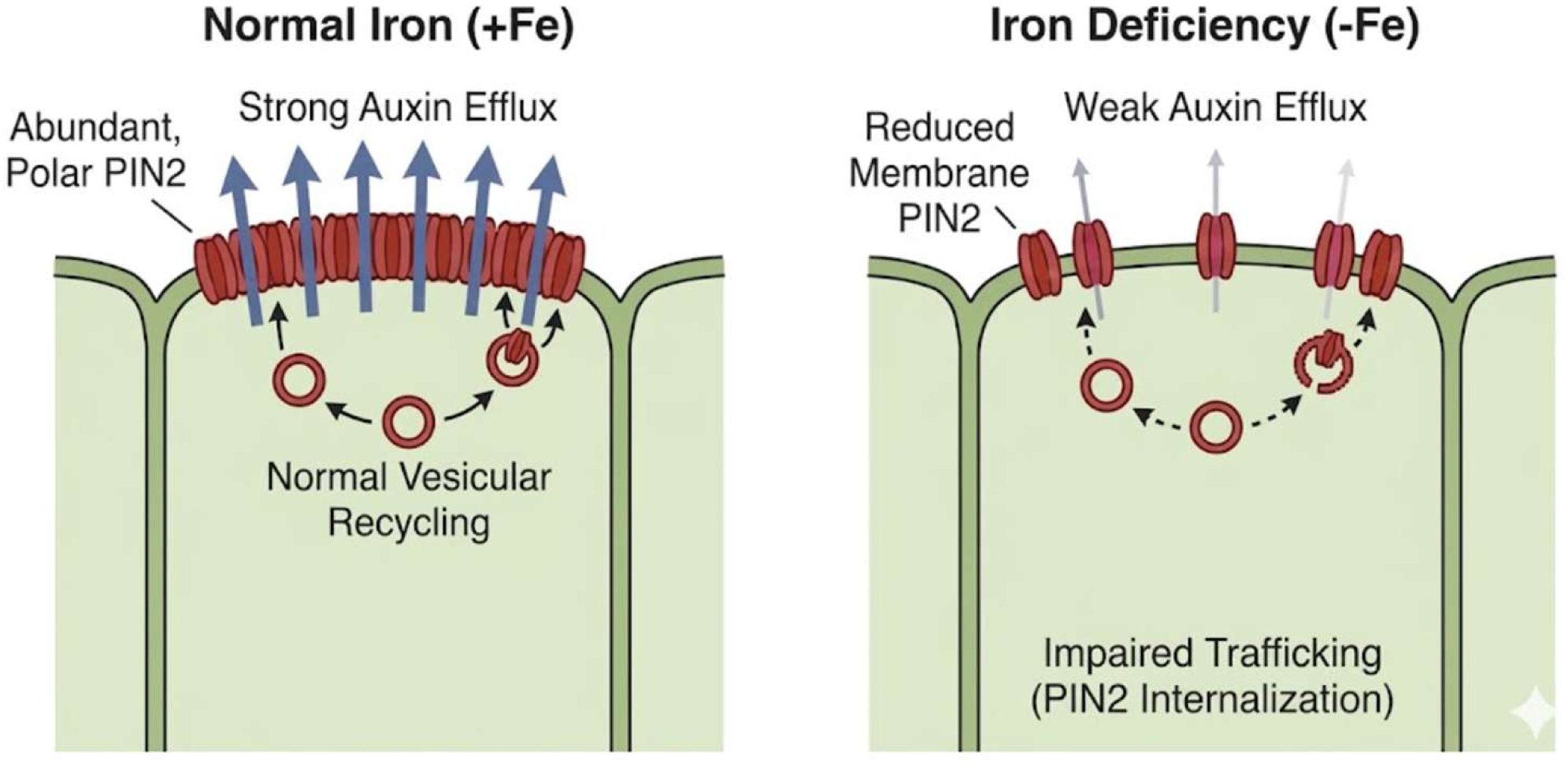
Hypothetical model of iron signaling affecting gravitropism via PIN2. Iron signaling mediates asymmetric auxin distribution at the root tip and affects gravitropism by regulating the abundance and localization of PIN2 protein. Under iron-sufficient conditions, PIN2 protein exhibits high abundance and polar localization at the plasma membrane, and intracellular vesicular trafficking remains normal, driving auxin efflux and basipetal transport, thereby ensuring asymmetric auxin distribution at the root tip and a normal gravitropic response. Under iron-deficient conditions, vesicular trafficking of PIN2 protein is impaired, its abundance at the plasma membrane is significantly reduced, and its polar localization is weakened; consequently, auxin efflux and basipetal transport are suppressed, disrupting asymmetric auxin distribution at the root tip and attenuating plant gravitropism.

Auxin plays a critical role in shaping the root system architecture, and multiple environmental cues have been reported to crosstalk with the auxin pathway to regulate RSA (Marchant et al., 1999; Overvoorde et al., 2010; Zhao, 2018; Szepesi, 2020; Tan et al., 2021). Here, our work indicates that iron regulates root gravitropic growth via modulating the subcellular localization of PIN auxin transporters and the subsequent auxin distribution. It would be interesting to further investigate which components involved in the auxin pathway requires iron and this would advance our understanding about the auxin-central network for root growth and development.

## 5 Conclusions

In summary, this study demonstrates that iron homeostasis mediated by the transporter IRT1 in Arabidopsis regulates asymmetric auxin distribution at the root apex and root gravitropic growth by selectively modulating the protein abundance, polar localization, and vesicle trafficking of the auxin efflux carrier PIN2. These results underscore the pivotal regulatory role of iron nutritional signals in root developmental plasticity, establishing a robust link between mineral nutrient sensing and auxin-mediated developmental programs.

Future research is required to further elucidate the molecular mechanisms by which iron signals directly intervene in endomembrane trafficking. Specifically, it remains to be determined whether iron-sensing proteins, such as the BRUTUS(BTS), directly mediate the post-translational modifications (PTMs) of PIN2, including ubiquitination or phosphorylation. Furthermore, a comprehensive analysis of how iron nutrition coordinates with other environmental cues and hormonal pathways will be essential for understanding how plants integrate multidimensional signals to optimize root system architecture (RSA) in response to fluctuating nutrient availability. Such efforts will facilitate the connection between nutrient transporters and hormonal signaling networks, revealing how mineral element homeostasis directly governs plant growth and development by orchestrating protein dynamics.

## Supporting Information

Supplemental information includes 14 Supplemental figures, 3 Supplemental tables and 4 Supplemental data sets.

## Acknowledgments

We thank Drs. Erika Isono (University of Constance), Grégory Vert (University of Toulouse), and Liwen Jiang (The Chinese University of Hong Kong) for kindly sharing published materials and the core facility of Life Sciences of the University of Science and Technology of China for their technical support. No conflicts of interest are declared. This work was supported by grants from the National Natural Science Foundation of China (32321001, 32570366 to S.T.), the Natural Science Foundation of Anhui Province (2508085QC070 to M.K.), the Fundamental Research Funds for the Central Universities (WK9100250095 to M.K., and WK9100000021 to S.T.), the Forestry Bureau of Anhui Province (AHLYJBGS-2024-01 to S.T.), the Center for Advanced Interdisciplinary Science and Biomedicine of IHM, Division of Life Sciences and Medicine, University of Science and Technology of China (QYPY20220012 to S.T.), the USTC Research Funds of the Double First-Class Initiative (YD9100002016 to S.T.), and start-up funding from the University of Science and Technology of China and the Chinese Academy of Sciences (GG9100007007, KY9100000026, KY9100000051, XKTS-202591014, and KJ2070000079 to S.T.).

## Preprint Statement

### Biographies

**Yao Fang** is currently a master’s student in the Division of Life Sciences and Medicine, University of Science and Technology of China, under the supervision of Prof. Shutang Tan. Her research direction is biochemistry and molecular biology.

**Yakun Peng** is currently a doctor in the Division of Life Sciences and Medicine, University of Science and Technology of China, under the supervision of Prof. Shutang Tan. Her research direction is biochemistry and molecular biology.

**Shutang Tan** is a professor at the Division of Life Sciences and Medicine, University of Science and Technology of China. His research interests include auxin biology, salicylic acid and plant development, plant cell and developmental biology and reversible protein phosphorylation.

## Author Contributions

S.T. conceived the project and designed the experiments. Y. F. and Y.P. performed the physiological and cell biological experiments under the supervision of S.T. All the authors contributed to the data analysis. Y. F., Y.P., and S.T. wrote the manuscript with input from other coauthors, and all authors revised and approved the submitted version.

