## supplemental Files for "Iron availability regulates PIN-mediated auxin transport and distribution to modulate root gravitropic growth in *Arabidopsis*"

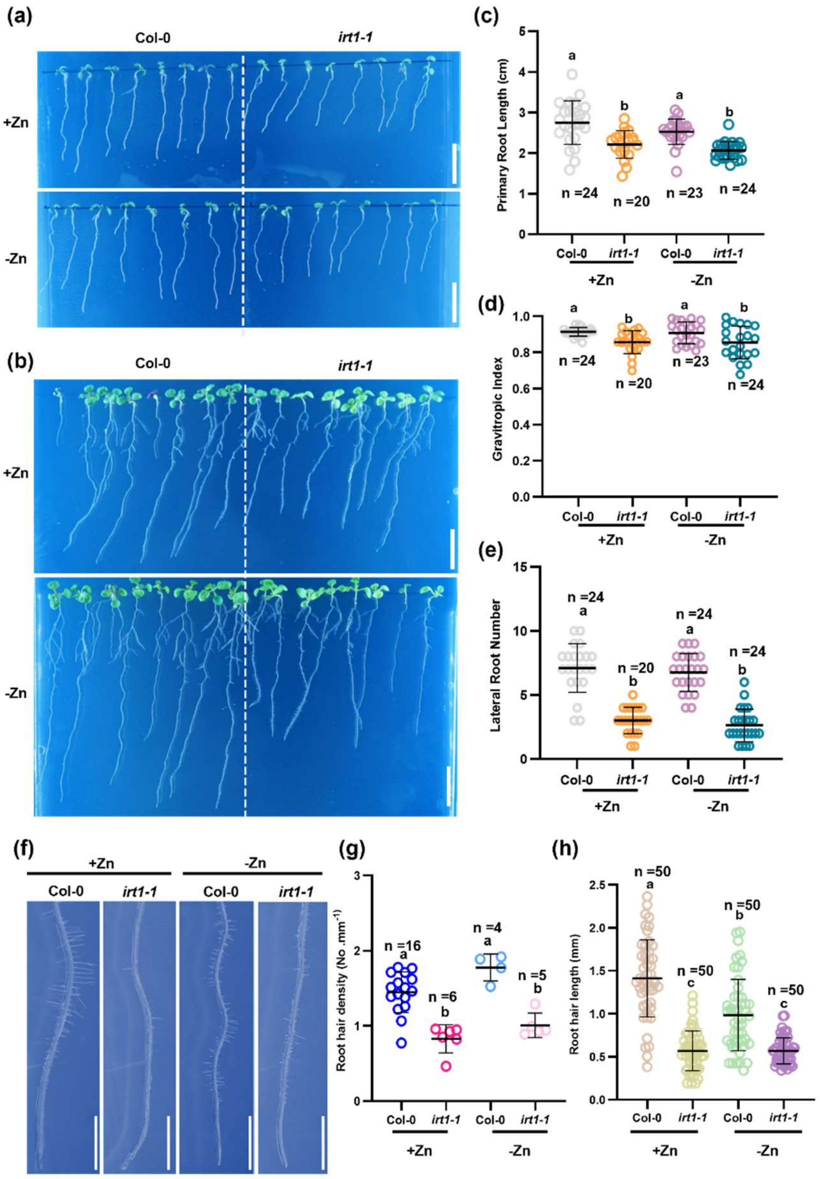


**Fig. S1. Zinc deficiency does not affect the gravitropism of Arabidopsis thaliana**

(a, b) Representative images of wild-type (Col-0) and *irt1-1* mutant seedlings grown for 7 and 11 days on MS media supplemented with zinc-sufficient(+Zn) or zinc-deficient(-Zn). Scale bars, 1 cm.

(c) Quantitative analysis of primary root length in *irt1-1* compared with Col-0 under normal and zinc-deficient conditions. *IRT1* deficiency resulted shorter roots; however, zinc deficiency did not further exacerbate this defect. The data are presented as the means ± SDs. Different letters denote statistically significant differences ($p < 0.05$, ANOVA with Tukey's HSD test).

(d) *IRT1* deficiency reduced gravitropism index(GI), with *irt1-1* mutants exhibiting gravitropic defects. Zinc deficiency did not further reduce the GI, nor did it affect gravitropism in Col-0 and *irt1-1*. Measurements were taken on seven-day-old Col-0 and *irt1-1* seedlings under normal and zinc-deficient conditions. The data are presented as the means ± SDs. Different letters denote significant differences; $P < 0.05$; one-way ANOVA with multiple comparisons.

(e) Quantitative analysis of the number of lateral roots in *irt1-1* mutants compared with Col-0 under normal and zinc-deficient conditions; the number of lateral roots in eleven-day-old and *irt1-1* seedlings was measured. *IRT1* deficiency disrupted normal lateral root development; the data are presented as the means ± SDs.

(f) *IRT1* deficiency inhibited root hair development, whereas zinc deficiency did not suppress normal root hair growth in Col-0. Root hair initiation was observed in elven-day-old Col-0 and *irt1-1* seedlings under various zinc conditions; the experiments were replicated five times, each yielding consistent results. Scale bars, 2.5 mm.

(g) Quantification of root hair density as shown. Data are presented as means ± SDs. Different letters indicate significant differences among groups by one-way ANOVA ($P < 0.05$).

(h) The figure illustrates the root hair length of Col-0 and *irt1-1* under zinc-sufficient(+Zn) and zinc-deficient(-Zn) conditions. Each circle represents an individual measurement from independent biological replicates ($n = 50$). Data are presented as means ± SDs. Different letters indicate significant differences among groups by one-way ANOVA ($P < 0.05$).

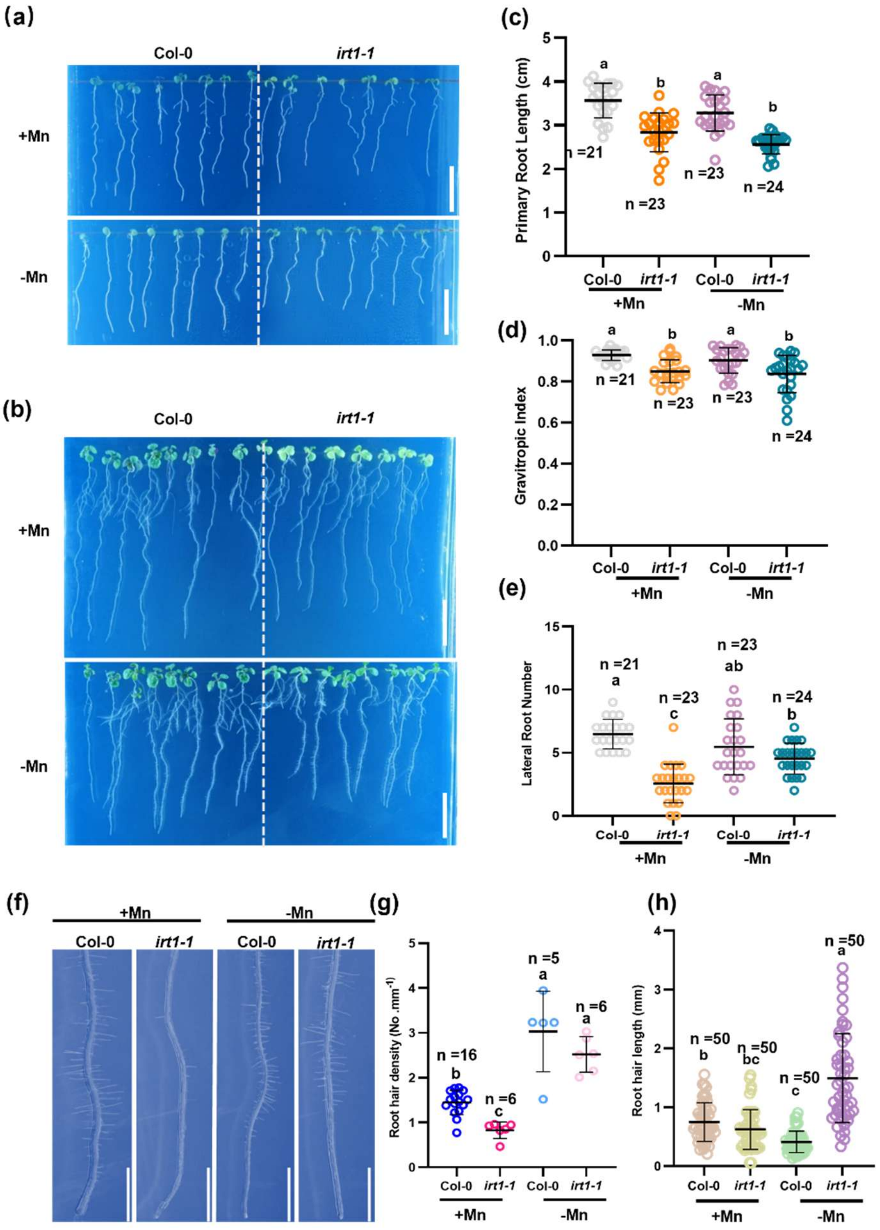


**Fig. S2. Manganese deficiency does not affect the gravitropism of Arabidopsis thaliana**

(a, b) Representative images of wild-type (Col-0) and *irt1-1* mutant seedlings grown for 7 and 11 days on MS media supplemented with manganese-sufficient(+Mn) and manganese-deficient (-Mn) conditions. Scale bars, 1 cm.

(c) Quantitative analysis of primary root length in *irt1-1* mutants compared with Col-0 under normal and manganese-deficient conditions. *IRT1* deficiency results in shorter roots; however, manganese deficiency did not further exacerbate this defect. The data are presented as the means ± SDs. Different letters denote statistically significant differences ($p < 0.05$, ANOVA with Tukey's HSD test).

(d) *IRT1* deficiency reduced gravitropism (GI), with *irt1-1* mutants exhibiting impaired gravitropism. Moreover, manganese deficiency did not further reduce the GI, and manganese deficiency did not affect the gravitropism of Col-0 or *irt1-1*. Measurements were taken using seven-day-old Col-0 and *irt1-1* seedlings under normal and manganese-deficient conditions. The data are presented as the means ± SDs. Different letters indicate significant differences; $P < 0.05$; one-way ANOVA with multiple comparisons.

(e) Quantitative analysis of the number of lateral roots of *irt1-1* and Col-0 under normal and manganese-deficient conditions. Lateral root numbers were measured in 11-day-old seedlings. The *IRT1* deficiency disrupted normal lateral root development. The data are presented as the means ± SDs.

(f) *IRT1* deficiency resulted in suppressed root hair development, whereas manganese deficiency did not inhibit normal root hair growth in Col-0. Observation of root hair initiation in 7-day-old Col-0 and *irt1-1* seedlings under manganese-sufficient and manganese-deficient conditions. The experiments were replicated five times with consistent results. Scale bars, 2.5 mm.

(g) Quantification of root hair density as shown. Data are presented as means ± SDs. Different letters indicate significant differences among groups by one-way ANOVA ($P < 0.05$).

(h) The figure illustrates the root hair length of Col-0 and *irt1-1* under manganese-sufficient(+Mn) and manganese-deficient (-Mn) conditions. Each circle represents an individual measurement from independent biological replicates ($n = 50$). Data are presented as means ± SDs. Different letters indicate significant differences among groups by one-way ANOVA ($P < 0.05$).

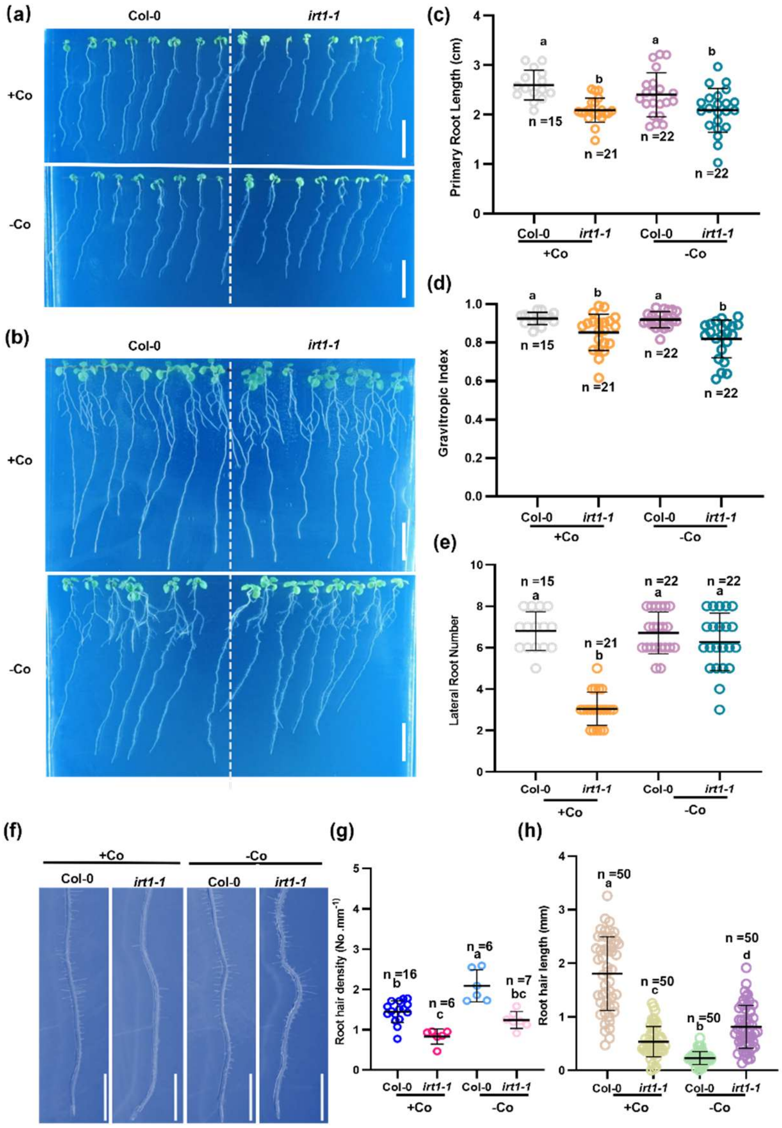


**Fig. S3. Cobalt deficiency does not affect the gravitropism of Arabidopsis thaliana**

**(**a,b) Representative images of Col-0 and *irt1-1* seedlings grown for 7 and 11 days on MS media supplemented with Co-sufficient (+Co) or Co-deficient (-Co) conditions. Scale bars, 1 cm.

(c) Quantitative analysis of primary root length in *irt1-1* compared with Col-0 under normal and cobalt-deficient conditions. *IRT1* deficiency resulted in shorter roots; cobalt deficiency did not further exacerbate this defect. The data are presented as the means ± SDs. Different letters denote statistically significant differences ($p < 0.05$, ANOVA with Tukey's HSD test).

(d) *IRT1* deficiency reduced gravitropism index(GI), with *irt1-1* exhibiting impaired gravitropism. Cobalt deficiency did not further reduce the GI, nor did it affect gravitropism in Col-0 or *irt1-1*. Measurements were taken using 7-day-old Col-0 and *irt1-1* seedlings under normal and cobalt-deficient conditions. The data are presented as the means ± SDs. Different letters denote significant differences; $P < 0.05$; one-way ANOVA with multiple comparisons.

(e) Quantitative analysis of lateral root number in *irt1-1* mutants versus wild-type Col-0 under normal and cobalt-deficient conditions. Lateral root numbers were measured in 11-day-old Col-0 and *irt1-1* seedlings. The *IRT1* mutation disruped normal lateral root development. The data are presented as the means ± SDs.

(f) *IRT1* deficiency inhibited root hair development, whereas cobalt deficiency did not suppress normal root hair growth in Col-0. Observation of root hair initiation in 7-day-old Col-0 and *irt1-1* seedlings under normal and cobalt-deficient conditions. The experiments were replicated five times with consistent results. Scale bars, 2.5 mm.

(g) Quantification of root hair density as shown. Data are presented as means ± SDs. Different letters indicate significant differences among groups by one-way ANOVA ($P < 0.05$).

(h) The figure illustrates the root hair length of Col-0 and *irt1-1* under Co-sufficient(+Co) and Co-deficient(-Co) conditions. Each circle represents an individual measurement from independent biological replicates ($n = 50$). Data are presented as means ± SDs. Different letters indicate significant differences among groups by one-way ANOVA ($P < 0.05$).

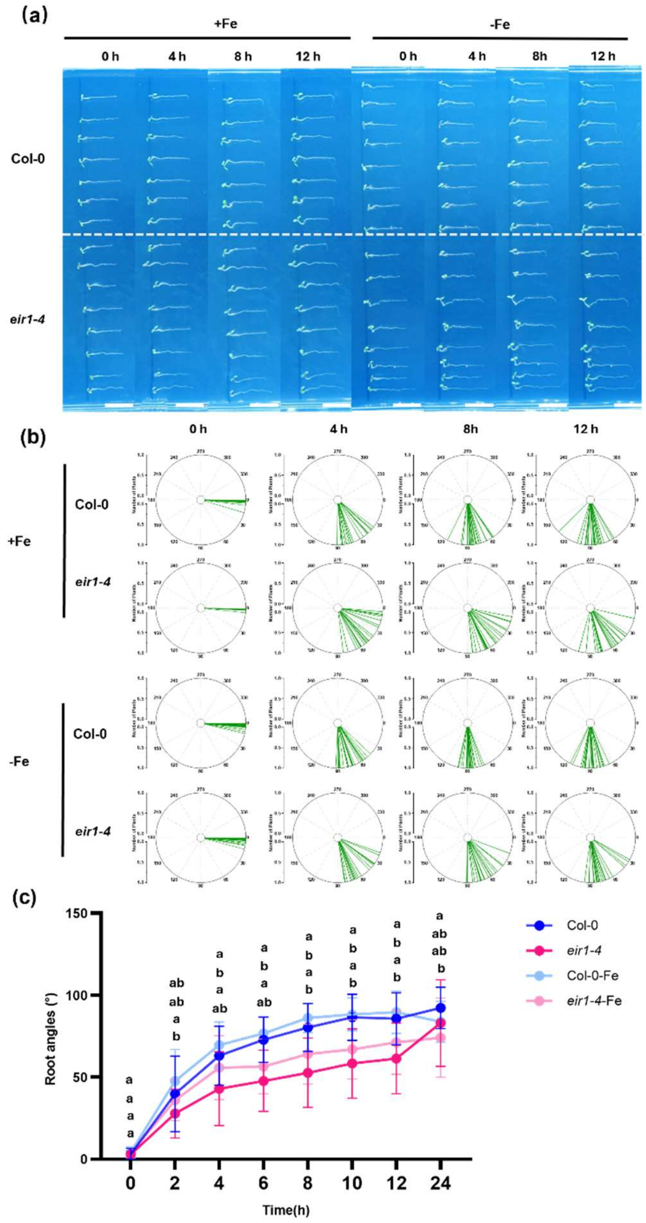


**Fig. S4. PIN2 deficiency disrupts normal root gravitropism in Arabidopsis**

(a) Representative root growth patterns at designated time points (0 h, 4 h, 8 h, 12 h) following a 90-degree rotation simulating gravitational stimulation after 4 days of growth on MS media under iron-sufficient (+Fe) and iron-deficient (-Fe) conditions in wild-type (Col-0) and *eir1-4* mutant seedlings. Scale bars, 1 cm.

(b) Root tip angles measured at 0 h, 4 h, 8 h, and 12 h post-gravistimulation in 4-day-old Col-0 and *irt1-1* seedlings under iron-sufficient (+Fe) and iron-deficient (-Fe) conditions, presented as polar histograms.

(c) Quantitative analysis of the root bending angles. At 2 h, 4 h, 6 h, 8 h, 10 h, 12 h, 24 h, the root bending angles of the *irt1-1* were significantly smaller than those of Col-0. The *irt1-1* mutant (irt1-1-Fe) exhibited impaired gravitropic responses, whereas Col-0 similarly displayed impaired gravitropic responses under iron deficiency. The data are presented as the means ± SDs.

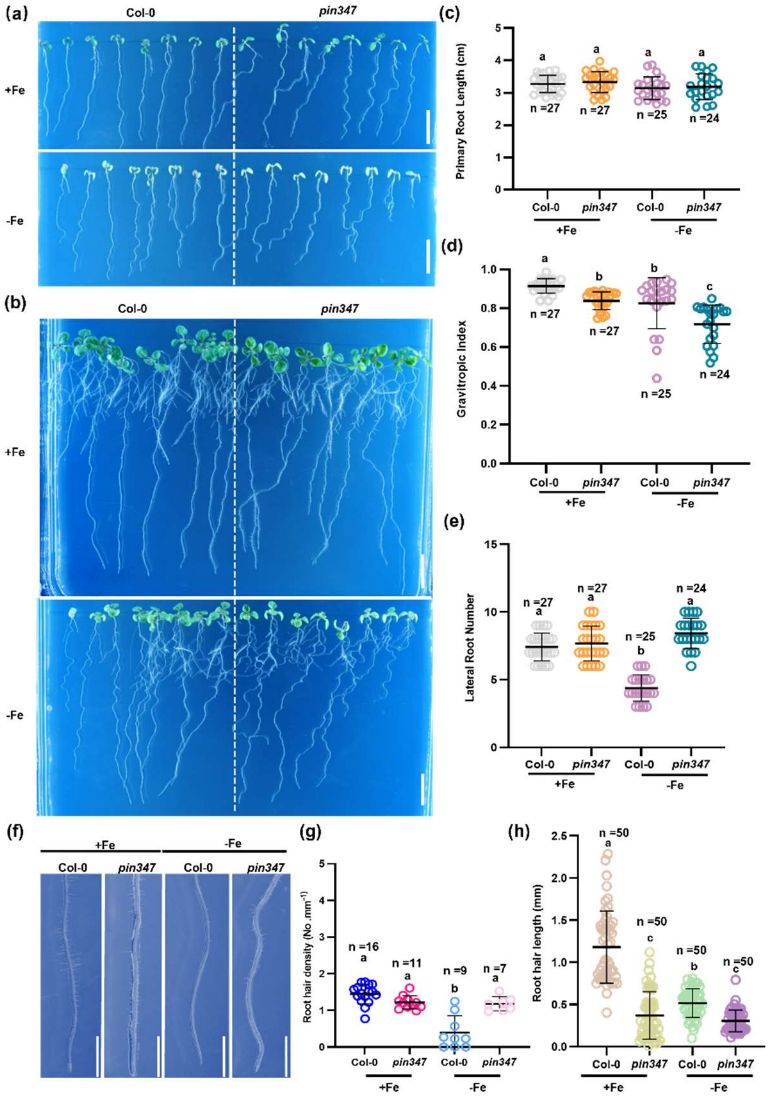


**Fig. S5. Iron deficiency further exacerbates the gravitropic defects of *pin347***

(a, b) Representative images of 7-day-old and 11-day-old Col-0 and pin347 seedlings grown on corresponding MS media under normal iron and iron-deficient conditions. Scale bars, 1 cm.

(c) Quantification of primary root length in *pin347* mutants compared with Col-0 under normal iron and iron-deficient conditions. The data are presented as the means ± SDs.

(d) The *pin347* mutant exhibited impaired gravity perception and a reduced GI index. Measurements were performed on 7-day-old Col-0 and *pin347* seedlings under normal iron and iron-deficient conditions. The data are presented as the means ± SDs. Different letters denote significant differences; $P < 0.05$; one-way ANOVA with multiple comparisons.

(e) Under iron deficiency, lateral root development is normal in both Col-0 and *pin347* mutants. Lateral root number was measured in 11-day-old seedlings. The data are presented as the means ± SDs.

(f) Iron deficiency inhibited normal root hair development in Col-0, whereas *pin347* mutants did not affect root hair formation. The initial stage of root hair emergence was observed in 7-day-old seedlings under various iron conditions. The experiments were performed in quadruplicate with consistent results. Scale bars, 2.5 mm.

(g) Quantification of root hair density as shown. Data are presented as means ± SDs. Different letters indicate significant differences among groups by one-way ANOVA ($P < 0.05$).

(h) The figure illustrates the root hair length of Col-0 and *pin347* under Fe-sufficient(+Fe) and Fe-deficient(-Fe) conditions. Each circle represents an individual measurement from independent biological replicates ($n = 50$). Data are presented as means ± SDs. Different letters indicate significant differences among groups by one-way ANOVA ($P < 0.05$).

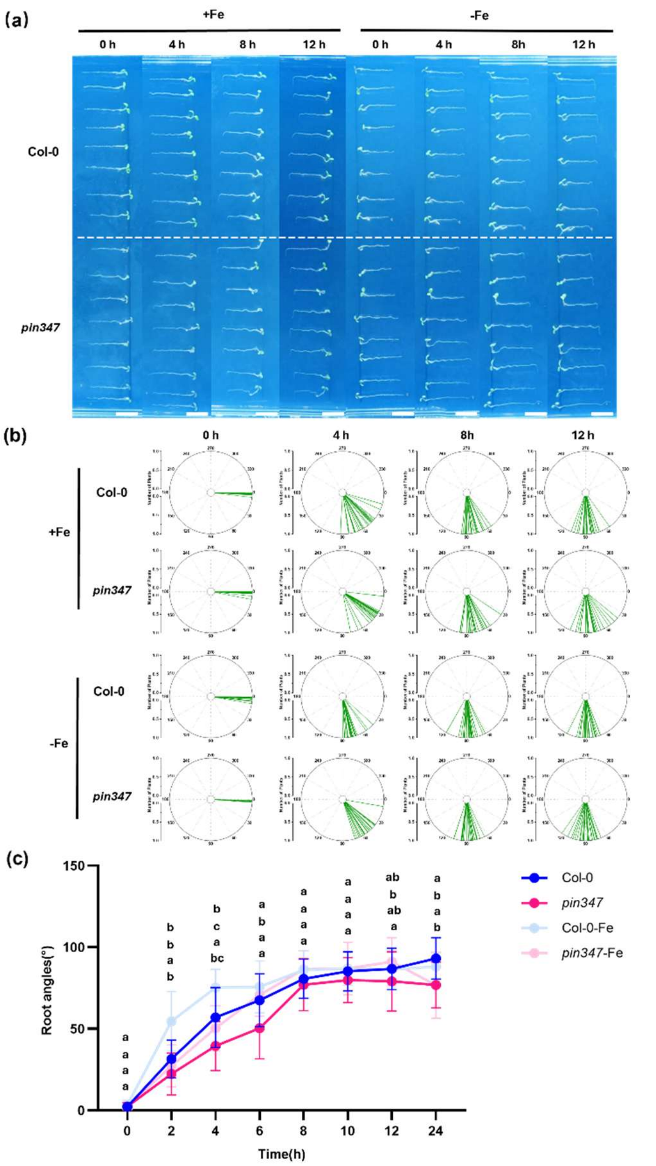


**Fig. S6. Defects in PIN3, PIN4, and PIN7 disrupt the normal gravitropic response of *Arabidopsis* roots**

(a) Root growth patterns of wild-type (Col-0) and *pin347* mutant seedlings after 4 days of growth on MS media under iron-sufficient (+Fe) and iron-deficient (-Fe) conditions, followed by 90° rotation to simulate gravitational stimulation. Images were captured at designated time points (0 h, 4 h, 8 h, and 12 h). Scale bars, 1 cm.

(b) The root tip angles of 4-day-old seedlings under iron-sufficient (+Fe) and iron-deficient (-Fe) conditions were measured at 0 h, 4 h, 8 h, and 12 h post-gravistimulation and presented as polar coordinate histograms.

(C) Quantitative analysis of root bending angles. At 2 h, 4 h, and 6 h, the root bending angles of the *pin347* mutant were smaller than those of Col-0 and converged at 8 h. The data are presented as the means ± SDs.

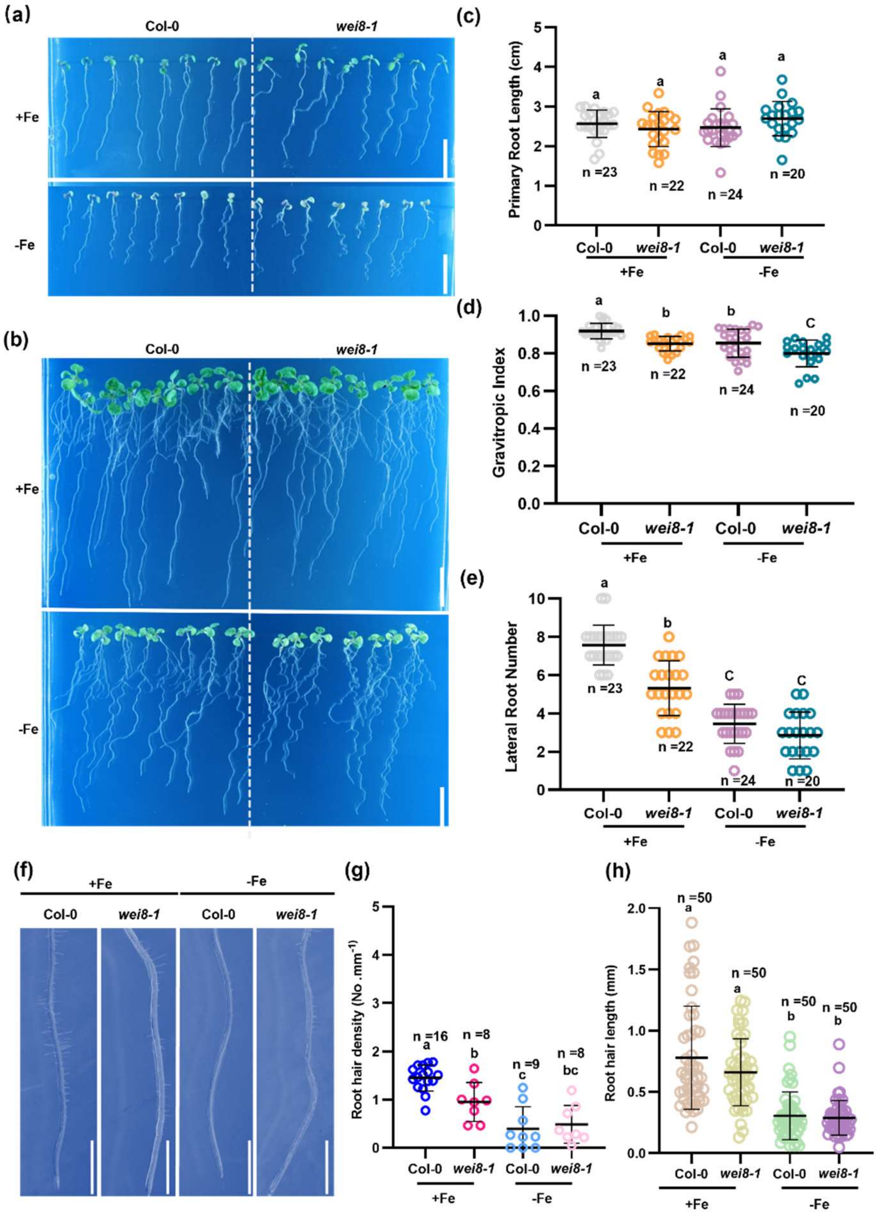


**Fig. S7. Iron deficiency affects the gravitropism of the *wei8-1* mutant**

(a,b) Representative images of 7-day-old and 11-day-old Col-0 and *wei8-1* seedlings grown on corresponding MS media under normal iron and iron-deficient conditions. Scale bars, 1 cm.

(c) Quantitative analysis of primary root length in the *wei8-1* mutant compared with Col-0 under normal iron and iron-deficient conditions. The data are presented as the means ± SDs.

(d) The *wei8-1* mutant exhibited impaired gravitropism, which affects the GI. Iron deficiency further impacted the GI, exacerbating gravitropic impairment, which was consistent with Col-0. Measurements were taken using 7-day-old Col-0 and *wei8-1* seedlings under normal iron and iron-deficient conditions. The data are presented as the means ± SDs. Different letters denote significant differences; $P < 0.05$; one-way ANOVA with multiple comparisons.

(e) Under iron deficiency, normal lateral root growth was inhibited in both Col-0 and *wei8-1*. The number of lateral roots was measured in 11-day-old seedlings. The data are presented as the means ± SDs.

(f) Compared with Col-0, root hair development in the *wei8-1* mutant is suppressed under normal iron conditions. Iron deficiency inhibited normal root hair development in Col-0, whereas root hair development in the *wei8-1* mutant remains unaffected. The experiment was replicated four times with consistent results. Scale bars, 2.5 mm.

(g) Quantification of root hair density as shown. Data are presented as means ± SDs. Different letters indicate significant differences among groups by one-way ANOVA ($P < 0.05$).

(h) The figure illustrates the root hair length of Col-0 and *wei8-1* under Fe-sufficient(+Fe) and Fe-deficient(-Fe) conditions. Each circle represents an individual measurement from independent biological replicates ($n = 50$). Data are presented as means ± SDs. Different letters indicate significant differences among groups by one-way ANOVA ($P < 0.05$).

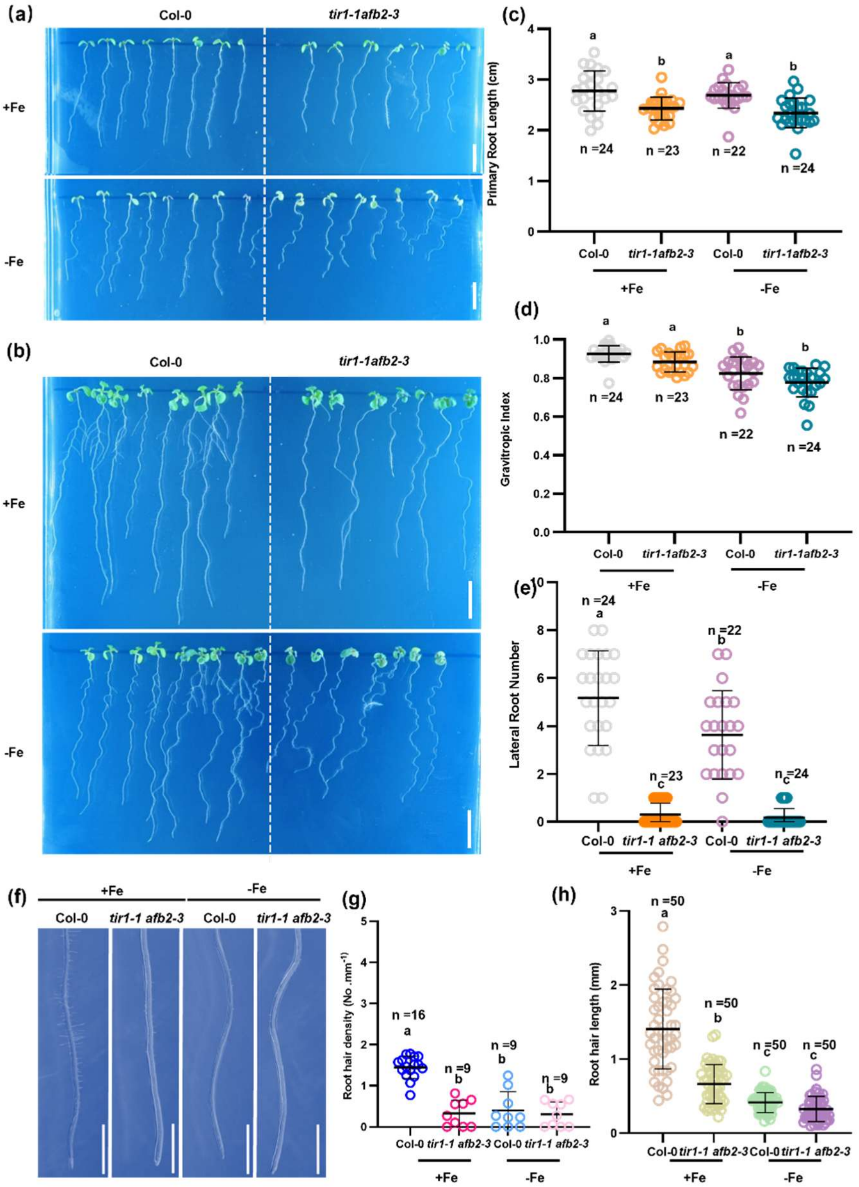


**Fig. S8. Iron deficiency enhances the gravitropic defects of the *tir1-1 afb2-3* mutant**

(a,b) Representative images of 7-day-old and 11-day-old Col-0 and *tir1-1 afb2-3* seedlings grown on corresponding MS media under normal iron and iron-deficient conditions. Scale bars, 1 cm.

(c) Quantitative analysis of primary root length in *tir1-1 afb2-3* mutants compared with Col-0 under normal iron and iron-deficient conditions. The data are presented as the means ± SDs.

(d) The *tir1-1 afb2-3* mutant exhibited reduced GI and impaired gravitropism. Iron deficiency further impacted the GI, exacerbating gravitropism impairment, which was consistent with the findings in Col-0. Measurements were taken from 7-day-old Col-0 and *tir1-1 afb2-3* seedlings under normal and iron-deficient conditions, and the data are presented as the means ± SDs. Different letters denote significant differences; $P < 0.05$; one-way ANOVA with multiple comparisons.

(e) Compared with Col-0, lateral root development in the *tir1-1 afb2-3* mutant was inhibited. Lateral root numbers were measured in 11-day-old Col-0 and *wei8-1* seedlings. The data are presented as the means ± SDs.

(f) Compared with Col-0, root hair development in the *tir1-1 afb2-3* mutant was suppressed under normal iron conditions. Iron deficiency inhibited normal root hair growth in Col-0, whereas root hair development in the *tir1-1 afb2-3* mutant remains unaffected under normal iron conditions and was completely suppressed. Observation of root hair initiation in 7-day-old Col-0 seedlings and *tir1-1 afb2-3* mutant seedlings under different iron conditions. The experiment was repeated four times with consistent results. Scale bars, 2.5 mm.

(g) Quantification of root hair density as shown. Data are presented as means ± SDs. Different letters indicate significant differences among groups by one-way ANOVA ($P < 0.05$).

(h) The figure illustrates the root hair length of Col-0 and *tir1-1 afb2-3* under Fe-sufficient(+Fe) and Fe-deficient(-Fe) conditions. Each circle represents an individual measurement from independent biological replicates ($n = 50$). Data are presented as means ± SDs. Different letters indicate significant differences among groups by one-way ANOVA ($P < 0.05$).

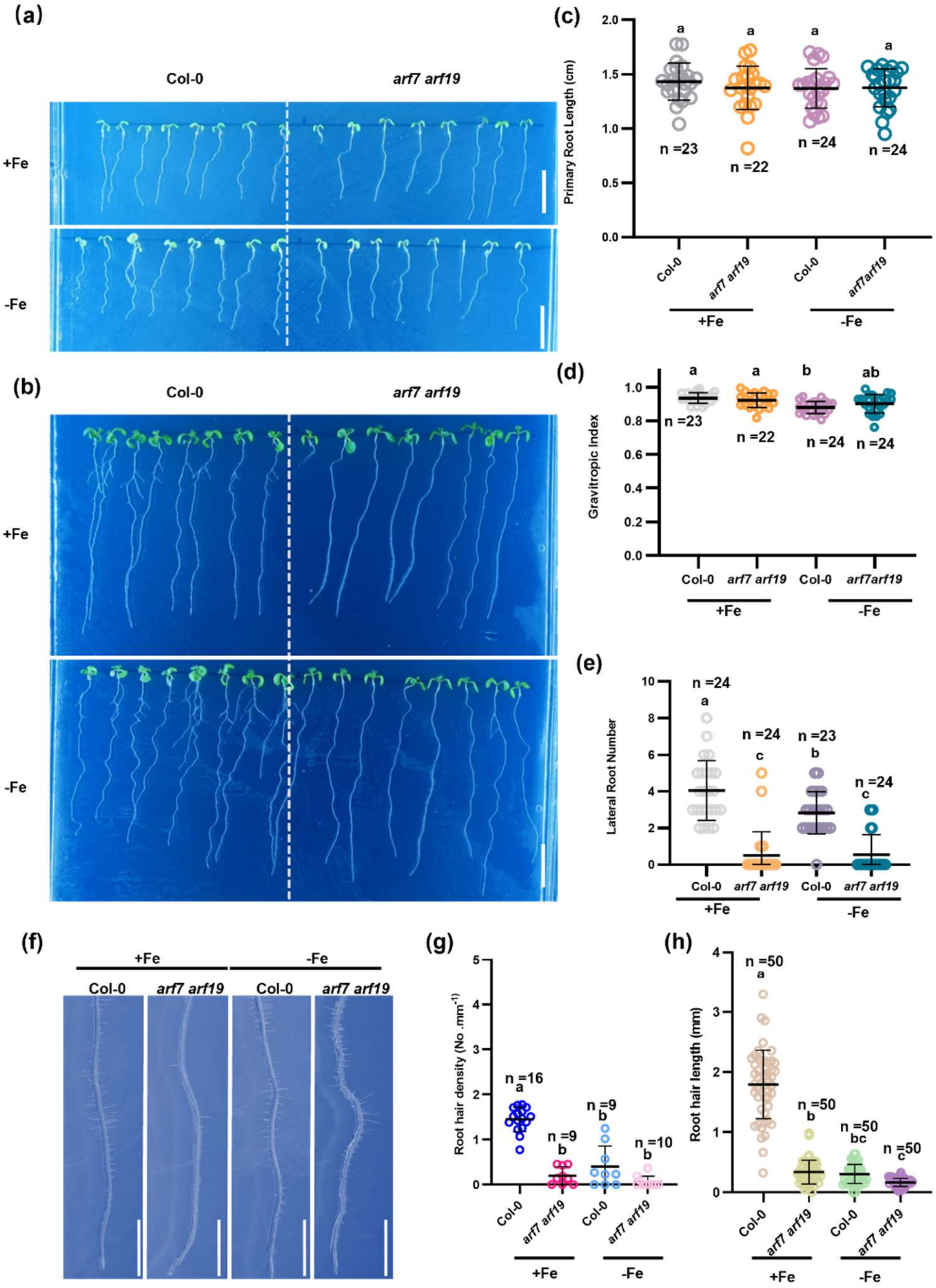


**Fig. S9. Iron deficiency affects lateral root growth in *arf7 arf19* mutants**

(a,b) Representative images of 7-day-old and 11-day-old Col-0 and *arf7 arf19* seedlings grown on corresponding MS media under normal iron and iron-deficient conditions. Scale bars, 1 cm.

(c) Quantitative analysis of primary root length in *arf7 arf19* mutants compared with Col-0 under normal iron and iron-deficient conditions. The data are presented as the means ± SDs.

(d) The GI of the *arf7 arf19* mutants were not significantly different from those of the Col-0. Measurements were taken using 7-day-old Col-0 and *arf7 arf19* seedlings under normal iron and iron-deficient conditions. The data are presented as the means ± SDs. Different letters denote significant differences; $P < 0.05$; one-way ANOVA with multiple comparisons.

(e) Compared with Col-0, normal lateral root development was significantly suppressed in *arf7 arf19* mutants. The number of lateral roots was measured in 11-day-old seedlings. The data are presented as the means ± SDs.

(f) Compared with Col-0, root hair growth in the *arf7 arf19* mutant was suppressed under normal iron conditions. Iron deficiency inhibited normal root hair growth in Col-0, whereas root hair development in the *arf7 arf19* mutant remained unaffected and was completely suppressed under normal iron conditions. The initial stages of root hair formation were observed in 7-day-old seedlings under various iron conditions. The experiment was replicated ten times, yielding consistent results. Scale bars, 2.5 mm.

(g) Quantification of root hair density as shown. Data are presented as means ± SDs. Different letters indicate significant differences among groups by one-way ANOVA ($P < 0.05$).

(h) The figure illustrates the root hair length of Col-0 and *arf7 arf19* under Fe-sufficient(+Fe) and Fe-deficient(-Fe) conditions. Each circle represents an individual measurement from independent biological replicates ($n = 50$). Data are presented as means ± SDs. Different letters indicate significant differences among groups by one-way ANOVA ($P < 0.05$).

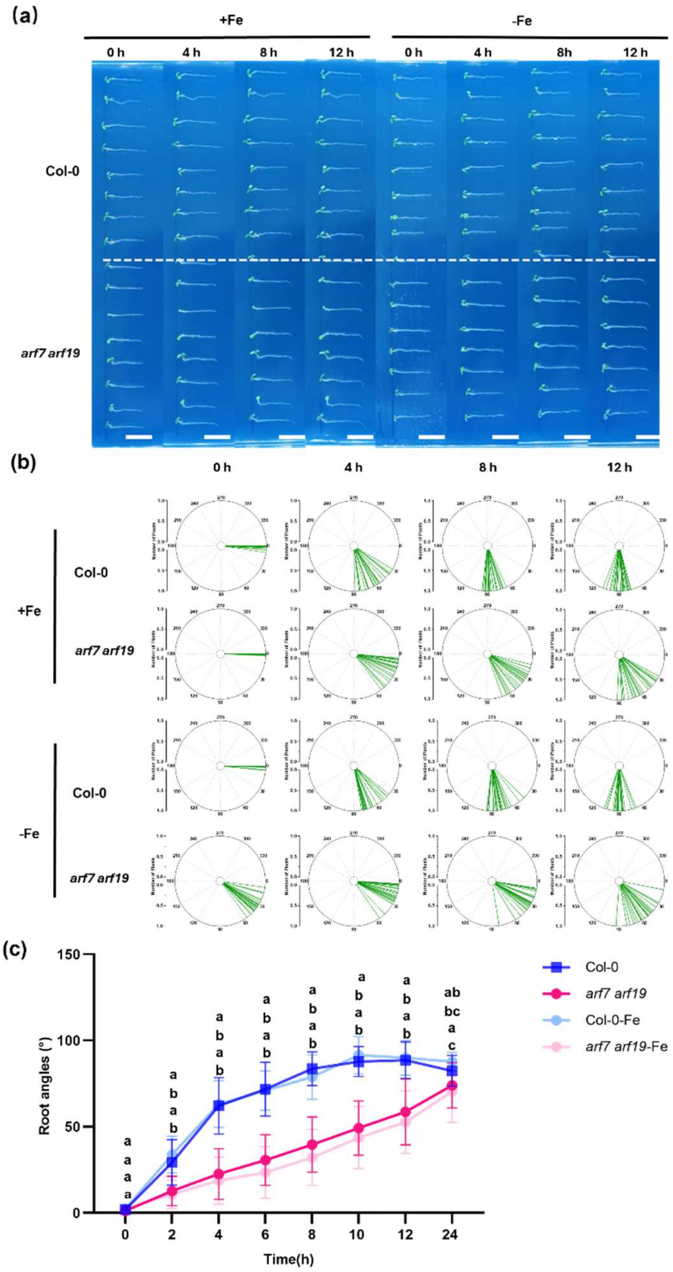


**Fig. S10. Gravitropic response of *arf7arf19* mutant seedlings**

(a) Representative root growth patterns of wild-type (Col-0) and *arf7 arf19* mutant seedlings after 4 days of growth on MS media under iron-sufficient (+Fe) and iron-deficient (-Fe) conditions, followed by 90° rotation to simulate gravitational stimulation. Images were captured at designated time points (0 h, 4 h, 8 h, and 12 h). Scale bars, 1 cm.

(b) The Root tip angles of 4-day-old Col-0 and *arf7 arf19* mutant seedlings under iron-sufficient (+Fe) and iron-deficient (-Fe) conditions, measured at 0 h, 4 h, 8 h, and 12 h post-gravitatory stimulation, presented as polar coordinate histograms.

(c) Quantitative analysis of the root curvature angle. At 2 h, 4 h, 6 h, 8 h, 10 h, and 12 h, the root curvature angle of the *tir1-1 afb2-3* mutant was significantly smaller than Col-0, converging towards parity at 24 hours. The data are presented as the means ± SDs.

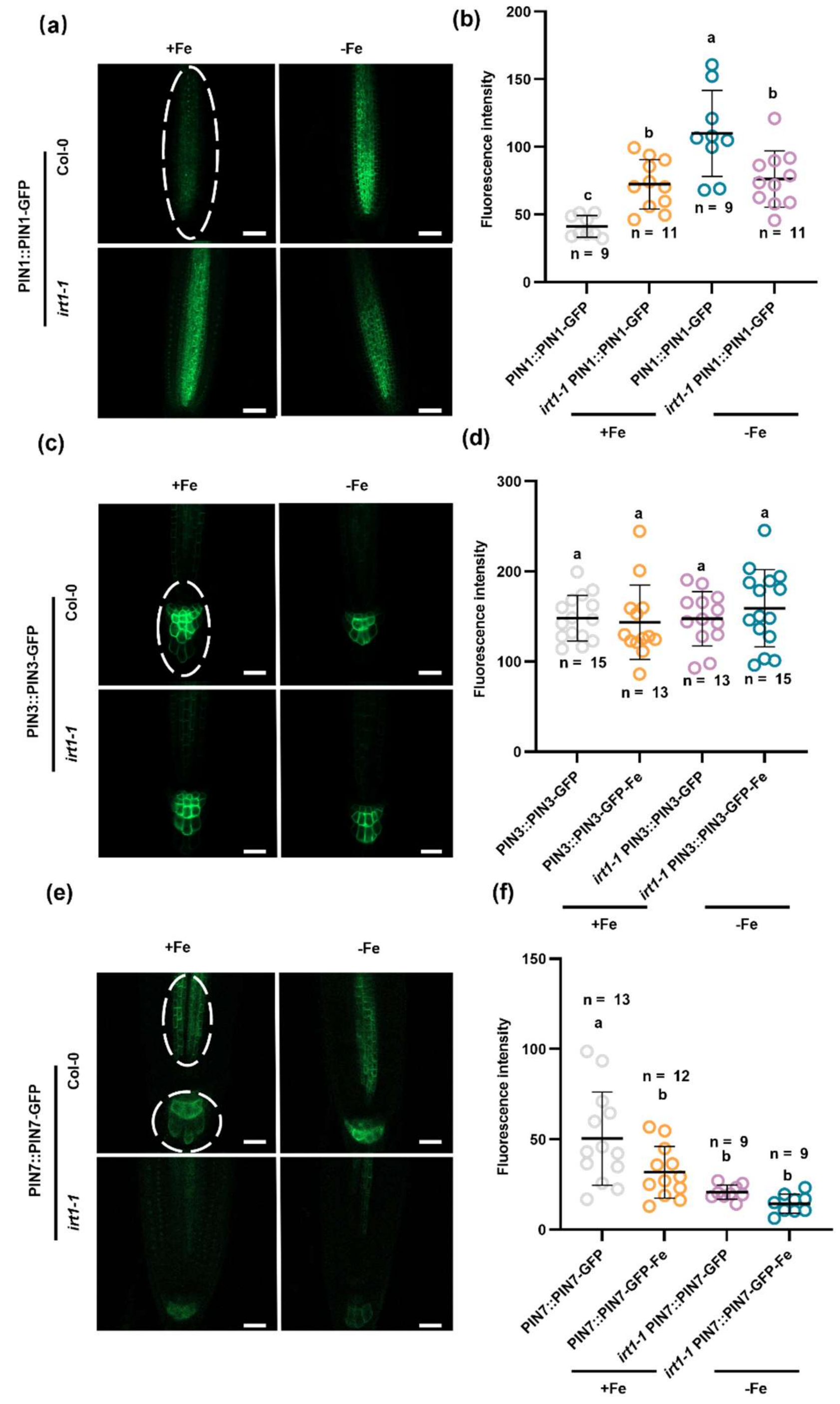


**Fig. S11. Iron deficiency and IRT1 deficiency interfere with the fluorescence of PIN1 and PIN7 but do not affect the fluorescence of PIN3**

(a) Representative images of *PIN1::PIN1-GFP* expression in *irt1-1* seedlings grown for 5 days on MS media with normal iron or iron deficiency; Col-0 was used as the control. Imaging parameters: laser wavelength, 488 nm, 5.00%; offset, 0.0%; detector gain, 700 V. Scale bar, 50 µm.

(b) In Col-0, the protein abundance of *PIN1::PIN1-GFP* was increased under iron deficiency. The fluorescence intensity of the same local area size was measured to assess reporter gene expression.

The data are presented as the means ± SDs; different letters indicate significant differences ($P < 0.05$); one-way analysis of variance (ANOVA) with multiple comparisons was used. The experiment was independently repeated three times, and similar results were obtained.

(c) Representative images of *PIN3::PIN3-GFP* expression in *irt1-1* seedlings grown for 5 days on MS media with normal iron or iron deficiency; Col-0 was used as the control. Imaging parameters: laser wavelength, 488 nm, 4.00%; offset, 0.0%; detector gain, 600 V. Scale bar, 20 μm.

(d) *PIN3::PIN3-GFP* expression in *irt1-1* seedlings resembled that in Col-0 plants, and iron deficiency did not affect its expression. The fluorescence intensity was measured at equal localised areas to demonstrate reporter expression; the data are presented as the means ± SDs; different letters denote significant differences; $P < 0.05$; one-way ANOVA for multiple comparisons. The experiments were independently replicated three times with similar results.

(e) Representative images of *PIN7::PIN7-GFP* expression in *irt1-1* seedlings grown for 5 days on normal iron and iron-deficient MS media, with Col-0 as a control. Epi-laser wavelength, 488 nm, 5.00%; 0.0% offset; detector gain, 800 V. Scale bar, 20 μm.

(f) Iron deficiency attenuated PIN7::PIN7-GFP expression in Col-0. *PIN7::PIN7-GFP* fluorescence was lower in the *irt1-1* seedlings than Col-0. The fluorescence intensity was measured across regions of equal size to visualise reporter expression. The data are presented as the means ± SDs; different letters denote significant differences; $P < 0.05$; one-way ANOVA for multiple comparisons. The experiments were independently replicated three times with similar results.

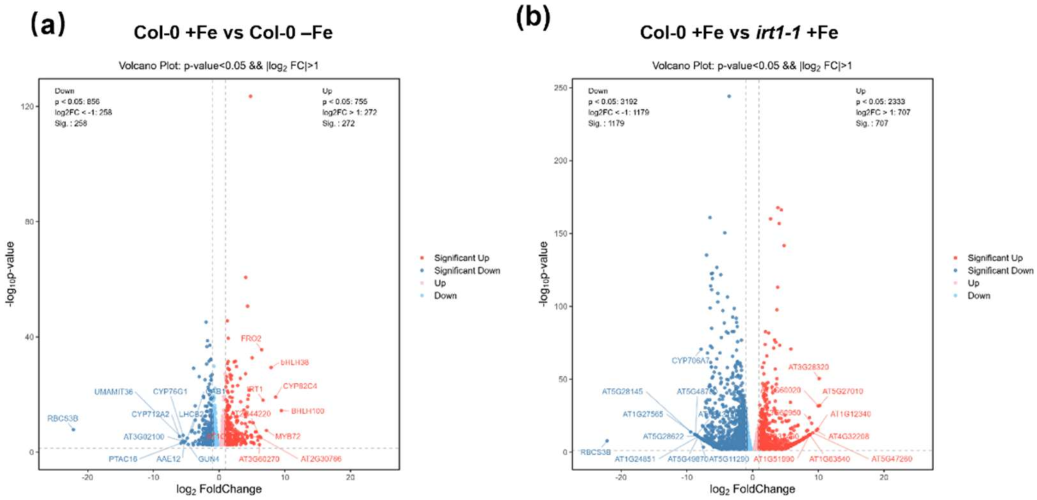


**Fig. S12. Volcano plots comparing gene expression profiles under different iron conditions and genetic backgrounds**

(a) Volcano plot comparing gene expression in Col-0 under iron-sufficient versus iron-deficient conditions (Col-0 +Fe vs Col-0 –Fe). Genes with significant differential expression (p value $\leq 0.05$ and $|\log_2 FC| \geq 1$) are highlighted as either significantly upregulated (red) or downregulated (blue). Nonsignificant genes are shown in gray. Select gene identifiers are annotated.

(b) Volcano plot comparing gene expression in Col-0 under iron-sufficient conditions versus *irt1-1* mutant plants under iron-sufficient conditions (Col-0 +Fe vs *irt1-1* +Fe). The same significance thresholds and color scheme as in (A) were applied. Both plots illustrate the overall distribution of $\log_2$-fold change versus statistical significance ($-\log_{10}$ p value). Key significantly upregulated and downregulated genes are labelled with their identifiers. The horizontal dashed line indicates the p value threshold (0.05), and the vertical dashed lines mark the $|\log_2 FC| = 1$ thresholds.

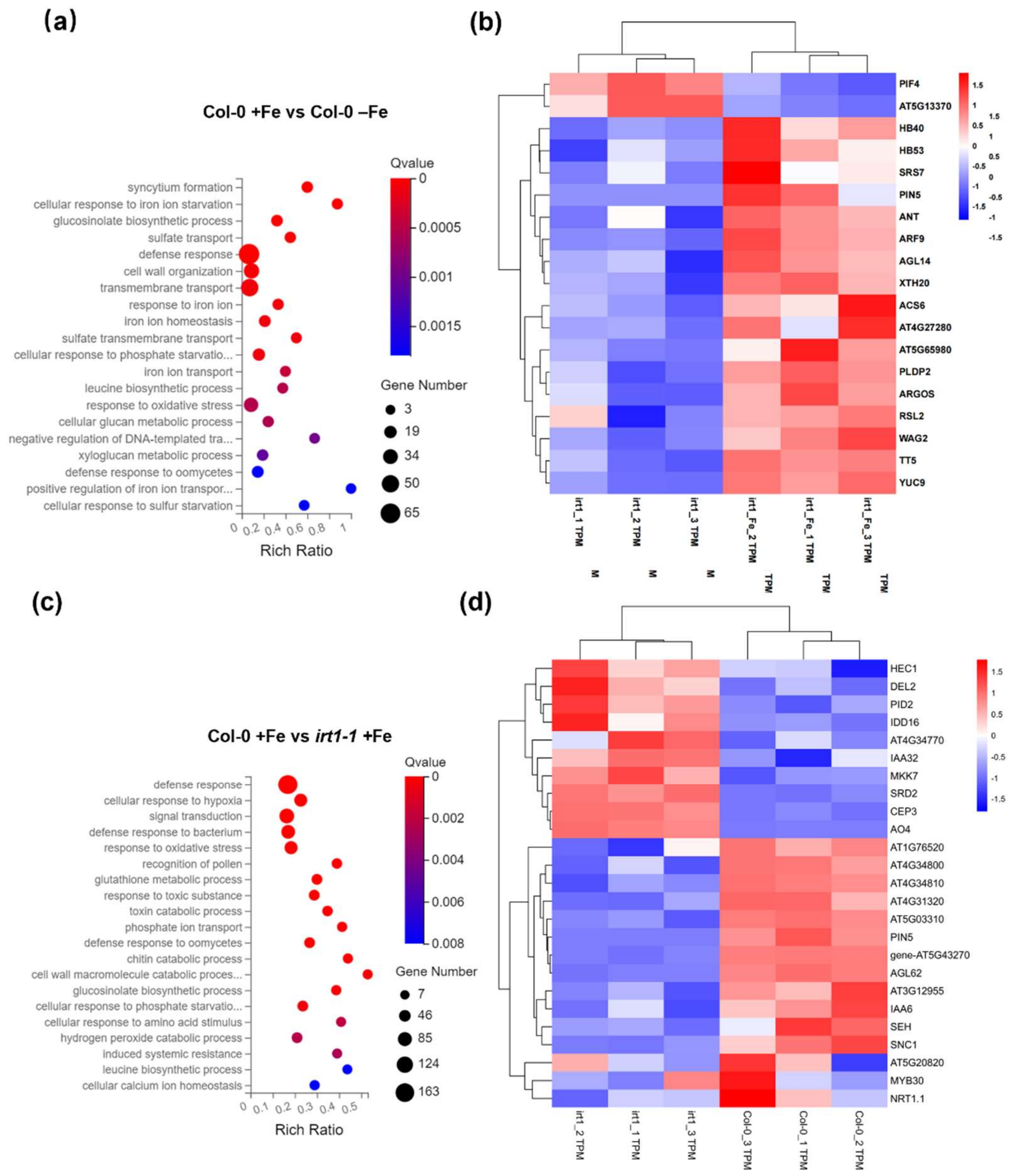


**Fig. S13. The RNA-seq results revealed altered gene expression in wild-type Col-0 and the *irt1-1* mutant following iron treatment**

(a) Statistics for GO enrichment analysis of differentially expressed genes in Col-0 under normal iron conditions versus *irt1-1* under iron deficiency.

(b) Heatmap displaying differentially expressed genes in Col-0 under normal iron conditions versus *irt1-1* under iron deficiency.

(c) Statistics from the GO enrichment analysis of genes differentially expressed between Col-0 and *irt1-1* plants under normal iron conditions.

(d) Heatmap displaying differentially expressed genes between Col-0 and *irt1-1* plants under normal iron conditions.

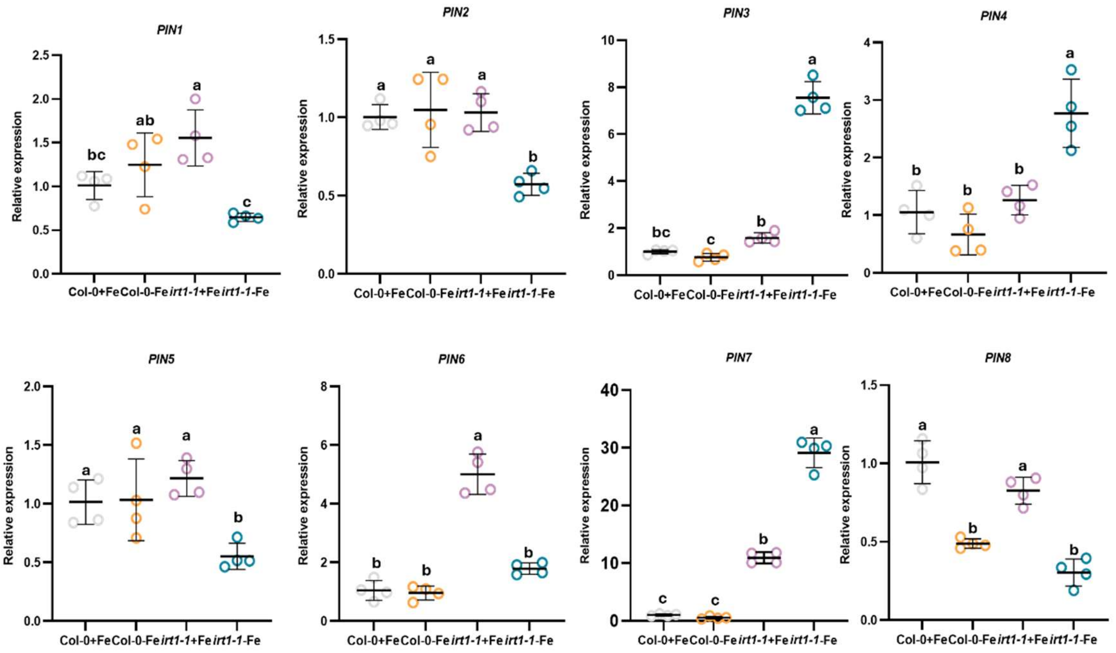


**Fig. S14. RT–qPCR analysis revealed the relative expression levels of PIN1–PIN8**

The dots represent individual data points (Col-0 in silver, *irt1-1* in yellow), while the lines denote the mean values ± standard deviations. Different letters denote statistically significant differences ($p < 0.05$, ANOVA with Tukey's HSD test).

**Supplementary Table 1. List of plant lines, including mutants and maker lines, used in this study.**

| *Arabidopsis* lines | Source | Identifier |
|---|---|---|
| Col-0 | N/A | N/A |
| *irt1-1* | (Vert et al., 2002) | N/A |
| *pPIN1::PIN1-GFP* | (Benková et al., 2003) | N/A |
| *pPIN2::PIN2-Venus* | (Retzer et al., 2019) | N/A |
| *pPIN3::PIN3-GFP* | (Žádníková et al., 2010) | N/A |
| *pPIN7::PIN7-GFP* | (Žádníková et al., 2010) | N/A |
| *DR5rev::GFP* | (Friml et al., 2003) | N/A |
| *arf7 arf19* | (Okushima et al., 2007) | N/A |
| *tir1-1 afb2-3* | (Ruegger et al., 1998) | N/A |
| *pin3 pin4 pin7* | (Blilou et al., 2005) | N/A |
| *wei8-1* | (Stepanova et al., 2008) | CS31113-19 |
| *eir1-4* | (Luschnig et al., 1998) | SALK_091142 |
| *irt1-1 pPIN1::PIN1-GFP* | This study | N/A |
| *irt1-1 pPIN2::PIN2-Venus* | This study | N/A |
| *irt1-1 pPIN3::PIN3-GFP* | This study | N/A |
| *irt1-1 pPIN7::PIN7-GFP* | This study | N/A |
| *irt1-1 DR5rev::GFP* | This study | N/A |

**Supplementary Table 2. List of reagents used in this study.**

| Reagent or Resource | Source | Identifier |
|---|---|---|
| **MS Base Salts with Vitamins** | Coolaber | Cat. # PM1011-100L |
| **MS Base Salts(-Fe,with vitamins)** | Coolaber | Cat. # PM1011-Fe-250G |
| **MS Base Salts(-Mn,with vitamins)** | Coolaber | Cat. # PM1011-Fe-250g |
| **1/2MS Base Salts(-Zn,with vitamins)** | Coolaber | Cat. # PM1061-Zn-250g |
| **MS Base Salts(-Co,with vitamins)** | Coolaber | Cat. # PM1011-Co-250g |
| **Plant Agar** | Duchefa | Cat. # P1001.1000 |
| **brefeldin A** | Beyotime | Cat. # S1536 |
| **Kits** | | |
| **GeneJET Plasmid Miniprep Kit** | Thermo Fisher Scientific | Cat. # K0503 |
| **FastPure Universal Plant Total RNA Isolation Kit** | Vazyme | Cat. # RC411-01 |
| **PrimeScript™ RT Master Mix (Perfect Real Time)** | Takara | Cat. # RR036A |
| **2 × Q3 Probe qPCR Master Mix** | Tolobio | Cat. # 22205 |
| **Software and Algorithms** | | |
| ***Arabidopsis* Information Resource (TAIR)** | N/A | https://www.arabidopsis.org/ |
| **Oebiotech** | N/A | https://cloud.oebiotech.cn/task/?id=19 |
| **BGI Biosystems** | N/A | https://biosys.bgi.com/#/report/project/projectList |
| **image J** | image J | https://imagej.nih.gov/ij/ |
| **Fiji** | Fiji | https://fiji.sc/ |
| **ZEN** | ZEN | https://www.zeiss.com/microscopy/en/products/software/zeiss-zen-lite.html |

**Supplementary Table 3. List of primers used in this study.**

| **Primers** | **Oligonucleotide (5' to 3')** | **Purpose** |
|---|---|---|
| **For Genotyping T-DNA Insertional Mutants** | | |
| **Oligo Name** | **Sequence** | **Allele** |
| *irt1-1*-LP | TAAAAGAAGATGATTCGTCAAATGCACAGC | *irt1-1* |
| *irt1-1*-RP | GTTAAGCCCATTTGGCGATAATCGACATT | *irt1-1* |
| *irt1-1*-LB | CTACAAATTGCCTTTTCTTATCGAC | *irt1-1* |
| **For qRT-PCR** | | |
| **Oligo Name** | **Sequence** | **Gene** |
| ACTIN7-F | CCGGTATTGTGCTCGATTCTG | *ACTIN7* |
| ACTIN7-R | TTCCCGTTCTGCGGTAGTGG | *ACTIN7* |
| CEP5-F | CATGGACGAACCCTAAAAGTTG | *CEP5* |
| CEP5-R | ATCTTCAGCATCTTTACCTCCC | *CEP5* |
| JAZ10-F | TCAGTTTTCCAAGTGTCTCGTA | *JAZ10* |
| JAZ10-R | TACCGAAAGATCTGTCTCCATC | *JAZ10* |
| ANT-F | GTAACACACTCTTGTCTGGAGA | *ANT* |
| ANT-R | GCCCATATTTGATCCGAACATC | *ANT* |
| MYB96-F | AGCTTCTTCCATGATCAAGTGA | *MYB96* |
| MYB96-R | CAAACAAAGACAGAGACCCTTG | *MYB96* |
| SAUR77-F | AGACCTTACATGCTTAGTTCCC | *SAUR77* |
| SAUR77-R | TTCGATAAATCTAGACGACCGG | *SAUR77* |
| MYB31-F | CCTCATCATCATTCTACCACCA | *MYB31* |
| MYB31-R | GAGACTGATGAAGTGTTAGGCT | *MYB31* |
| PIP2-F | CGTTCTGAGTTCGATTCTGTTC | *PIP2* |
| PIP2-R | ACTTCTCCTCGGTCTTTGTTAG | *PIP2* |
| GRDP2-F | TGAAAGTTTACTCCGGGAGAAA | *GRDP2* |
| GRDP2-R | CGAACTTCAAATCGAGTAACCC | *GRDP2* |
| IAA5-F | GACCAAAAGTTCGTACGTGAAA | *IAA5* |
| IAA5-R | CACATTCACTTTCCTTCAACGT | *IAA5* |
| IAA19-F | CGCTGAGAAGGTTAATGATTCG | *IAA19* |
| IAA19-R | TCACTTTCACATACCCTAACCC | *IAA19* |
| ARF7-F | TACCTGATGCAGCGATTGATAT | *ARF7* |
| ARF7-R | TTCTTGTTTTAGCCGCATATCG | *ARF7* |
| ARF9-F | CTGTTCTAAGGAAACATGCCAC | *ARF9* |
| ARF9-R | CCTCTCAGGAATACAAAGGTGT | *ARF9* |

| | | |
|---|---|---|
| LB29-F | ACAGAGAGTAGTTACCACAACG | *LB29* |
| LB29-R | CCTGATTGAAAGTGTTCAGGTG | *LB29* |
| GH3.3-F | CTGGCTGATTTCATAACTTCGG | *GH3.3* |
| GH3.3-R | CGATGACGTCAAGGTACTTAGT | *GH3.3* |
| PIN1-F | AGGGATGTTTTCGCCCAACA | *PIN1* |
| PIN1-R | AGTAATCGGCGTGGTGGTTT | *PIN1* |
| PIN2-F | TTCTTTGGCAGGCGTTTAGC | *PIN2* |
| PIN2-R | TAGCCCCACGGAACTCAAAC | *PIN2* |
| PIN3-F | TCAACCACCACATCTACCGC | *PIN3* |
| PIN3-R | TTCCGCCTTGATCGTTGTCA | *PIN3* |
| PIN4-F | CGGATTTGTACTCCGTTCAATC | *PIN4* |
| PIN4-R | GTTTAGTTGAAACACCCGTACC | *PIN4* |
| PIN5-F | TATCAGCGACGTACAAGTAGAC | *PIN5* |
| PIN5-R | AGAAATAAAAGCCCATGCGATC | *PIN5* |
| PIN6-F | GGAGATTACACTCAAACCCTCA | *PIN6* |
| PIN6-R | CATCGGTTTCAGTTTCTGTACG | *PIN6* |
| PIN7-F | AATTCGACGGCGACTTTTGC | *PIN7* |
| PIN7-R | TGAGGCAATGCAGCTTGAAC | *PIN7* |
| PIN8-F | CAAAGCTTGATTTGGTACACCA | *PIN8* |
| PIN8-R | ATTCCGATCAATGTTGCGTATG | *PIN8* |
